# Synovial fluid transcriptome dynamics in osteoarthritis progression: Implications in pathogenesis

**DOI:** 10.1101/2024.06.24.600143

**Authors:** Rinkle Sharma, Diksha Rana, Rahul Kumar, Sakshi Narula, Alpa Chaudhary, Bhavneet Kaur, Khushpreet Kaur, Mandeep Dhillon, Devendra K Chauhan, Uttam Chand Saini, Sadhna Sharma, Jyotdeep Kaur, Indu Verma

## Abstract

**Background:** Osteoarthritis, a degenerative joint disease associated with various pathological manifestations in the joint including cartilage loss, alterations in subchondral bone and synovial inflammation.

**Objective:** This study aimed to elucidate the transcriptional and molecular changes in synovial fluid associated with OA progression, focusing on differential gene expression and pathway enrichment across OA grades.

**Methodology:** Patients with different OA grades were recruited from PGIMER, Chandigarh, following the KL classification. Microarray analysis was conducted to study the transcriptional profiles in different OA grades using a fold-change (FC) cutoff of 2 and a p-value cutoff of 0.05, followed by pathway analysis performed using GSEA and STRING database. Selected genes from microarray and pathway analysis were validated using qRT-PCR.

**Results:** Microarray analysis reveals distinct gene expression patterns corresponding to different OA stages (KL grade 2 to KL grade 4). Notably, the upregulation of *AMTN* and *DKK2*, alongside the downregulation of *MSLN*, highlighted their roles in pathological mineralization and disrupted bone remodeling in OA. Pathway enrichment analysis revealed significant changes in immune response, inflammation related pathways and cellular processes such as autophagy and programmed cell death, indicating their involvement in disease progression. Furthermore, mitochondrial dysfunction and impaired autophagy were linked to increased inflammation in advanced OA.

**Conclusion:** These findings suggest that targeting mineralization and inflammatory pathways could offer novel therapeutic avenues for OA management.

## 1.0 Introduction

Osteoarthritis (OA) is the most debilitating and common form of arthritis. According to the World Health Organization, out of all rheumatologic problems, OA is the most common joint disease with a prevalence of 22% to 39% in India. It is likely that OA is triggered by advanced aging, high body weight, genetic predisposing factors, mechanical stress, and trauma. Clinically OA can affect any weight-bearing joint and is characterized by loss of cartilage, formation of bony spurs (osteophytes) margins of joints, subchondral bone sclerosis, synovial inflammation, and pathological mineralization (1, 2). The present lack of approved disease-modifying OA drugs (DMOADs) does not allow the prevention, halting or even limiting OA’s progression. As a result, OA is considered as an incurable and irreversible disease by the Osteoarthritis Research Society International (OARSI), however, its progression can only be staved off by treating it in time (3). The pathogenesis mechanism of OA has long been studied from the perspective of pathological changes in articular cartilage and chondrocytes but has not focused much on the inflammatory changes in the synovial fluid. Most physicians consider OA to be long-term cartilage loss and bone damage caused by degenerative cartilage degradation, particularly because OA is normally asymptomatic and does not accompany systemic inflammation. However, there is increasing evidence that synovial inflammation can produce inflammatory mediators that influence not only articular cartilage, but also some metabolic cytokines and proteases, thus accelerating joint deterioration (4, 5). Hence, it is imperative to have an insight into the various pathways associated with progression of OA disease particularly for better therapeutics to slow down the progression of the disease from its early stages into its late stages. Synovial fluid is the most affected biological fluid along with its proximity to all the affected tissues in OA joint (6). Analyzing the synovial fluid transcriptome is more advantageous for identifying potential biomarkers than studying serum or plasma. Since synovial fluid directly interacts with the synovium, ligaments, meniscus, joint capsule, and bone, it more accurately reflects molecular changes associated with disease in these tissues. Thus, examining synovial fluid can reveal critical insights into the severity and progression of osteoarthritis. This approach holds promise for early osteoarthritis diagnosis, monitoring disease progression, and informing treatment options.

The current study investigated the synovial fluid transcriptomics in OA patients across different disease stages, from KL grade 2 to KL grade 4. In depth analysis of transcriptomic data has led to identification of hub genes and associated pathways important to maintain joint structure at different stages of OA disease and may serve as important for the development of novel therapeutics.

## 2.0 Methodology

### 2.1 Ethical clearance and patient recruitment

Patients with symptoms of knee pain diagnosed with OA undergoing platelet-rich plasma (PRP) therapy and total knee replacements (TKR) were included in the study. Synovial fluid samples were collected from OA patients at the Department of Orthopaedics, Nehru Hospital, at the Post Graduate Institute of Medical Education and Research (PGIMER), Chandigarh. The study was conducted with informed consent and was approved by the PGIMER’s Institute Ethics Committee under the ethical clearance number PGI/IEC/2017/16 and 67 patients with different OA grades were recruited in the study.

Synovial fluid samples from different grades of OA patients were collected on the basis of the patient diagnosis that was made according to criteria laid down by the Subcommittee of the American College of Rheumatology Diagnostic and Therapeutic Criteria Committee on Osteoarthritis (7). Grading of the severity of knee OA of each patient was assessed by Kellgren Lawrence (KL) grading scales based on plain radiographs (8). Patients were categorised into three categories based on the basis of KL grading considering osteophyte formation, joint space narrowing, sclerosis, and joint deformity: Grade 2, Grade 3 and Grade 4 **(Supplementary figure 1)**. The grading of the samples was done by a trained orthopaedician.

Inclusion criteria were symptomatic (age- and sex-matched) patients, 55-70 years of age with knee pain. Patients suffering from other musculoskeletal diseases, anti-tubercular drug, history of taking glucocorticoids, anti-epileptic drugs, knee injections of cortisone or hyaluronic acid, trauma, ligament injury were excluded. None of the patients had diabetes.

Synovial fluid was collected from affected knee of enrolled patients undergoing PRP therapy or TKR surgery by arthrocentesis procedure under aseptic condition by a trained orthopaedician **(Supplementary figure 1)**. The synovial fluid was immediately transported to the lab on ice for total RNA isolation.

### 2.2 RNA extraction

RNA from synovial fluid samples was isolated using the commercially available kit (Qiagen RNeasy Mini kit), as per the protocol manual with slight modifications. In brief, synovial fluid was diluted 1:1 with nuclease-free water (NFW) to reduce its viscosity and centrifuged at 10,000 Xg for 10 minutes at 4°C to recover the pellet and the rest of the procedure was done as per the manufacturer’s protocol. Total RNA was eluted in an appropriate volume of NFW supplied with the kit. The concentration and the quality (260/280 ratio) of RNA were accessed by NanoDrop (Thermo Scientific). The RNA integrity was accessed by Agilent Bioanalyzer 2100 to confirm the quality of RNA required for microarray. The samples with good RIN values were further processed for mRNA microarray **(Supplementary figure 2 and table 1).** Samples with blood contamination were excluded from the study **(Supplementary figure 1).**

**Table 1.** Clinical characteristics of different grades of osteoarthritis patients as shown in the demographics.

| S.No. | KL Grade | No. of subjects | Age (in years)* | No. of females | WOMAC score** | BMI** (kg/m<sup>2</sup>) |
| --- | --- | --- | --- | --- | --- | --- |
| 1. | Grade 2 | 19 | 52±7.73 | 13 (68.42%) | 56.3±19.12 | 26.79±5.79 |
| 2. | Grade 3 | 7 | 61±5.53 | 6 (60%) | 60.42±13.51 | 27.97±4.56 |
| 3. | Grade 4 | 36 | 65±7.91 | 25 (69.44%) | 62.15±13.88 | 28±4.49 |

\*Median ± SD, \*\*Mean ± SD, KL-Kellgren Lawrence, WOMAC- Western Ontario and McMaster Universities Osteoarthritis, BMI- Body Mass Index

### 2.3 Transcriptome profiling of synovial fluid samples

Using an Agilent platform (Design ID: G4858A) commercial service, microarray was carried out to screen the differentially expressed genes (mRNA). For the mRNA microarray, input RNA (200ng) from the control and test samples was amplified using T7 RNA polymerase and labelled with Cyanine 3-CTP at the same time using the One-Color Microarray-Based Low Input QuickAmp Labelling Kit (Agilent Technologies). The Cyanine 3-labeled cRNA was quantified via spectrophotometry using a Nanodrop and purified using the RNeasy Mini Kit. The yield and specific activity were measured. The samples with a specific activity of more than 10 pmol were used for hybridization. The cyanine 3-labeled cRNA samples were hybridized to Agilent whole human genome 8*60k microarray chips. Following hybridization, the array slides were washed and scanned with an Agilent SureScan High-Resolution DNA Microarray Scanner. Feature extraction was performed using Feature Extraction Software, and the resulting signal intensities were analyzed using Agilent GeneSpring GX software version 14.9.1. The data was normalized through quantile normalization. To identify significantly differentially expressed entities among synovial fluid samples from varying grades of osteoarthritis (OA) patients, analysis of variance (ANOVA) was utilized, followed by a post-hoc Tukey test for intergrade comparisons. Entities with a p-value ≤ 0.05 and fold change ≤ -1.5 or ≥ 1.5 were considered significant.

### 2.4 Computational/ i*n silico* analysis

#### Pathway analysis

Gene Set Enrichment Analysis (GSEA) were used for enrichment analyses to investigate DEGs at the molecular and functional levels. In order to study the effect of the DEGs on various biological, molecular and cellular functions, GSEA (version 4.1.0, http://software.broadinstitute.org/gsea/downloads.jsp) was used to analyze DEGs of biological pathway annotation, cellular component annotation, molecular function and REACTOME annotation. The number of permutations was set at 1000, P value < 0.05 and a false discovery rate (FDR)<0.1 were considered statistically significant for enrichment results (9). Interactive relationships among DEGs were selected with a score (high confidence) >0.7, which was thought to be statistically significant. Search Tool for the Retrieval of Interacting Genes database (STRING version.11.5; http://string-db.org) was also used to study the network/protein-protein interaction (PPI) of their gene counterparts (10).

### 2.5 Validation using quantitative reverse transcription PCR (RT-qPCR)

Based on the data for transcriptomic analysis, selected genes were further validated in the synovial fluid samples of study subjects by real-time RT-qPCR. Total RNA was isolated from each sample (synovial fluid) and was processed with the First Strand cDNA Synthesis Kit according to the manufacturer’s protocol (BioRad iScript kit). Primers were synthesized from sigma for shortlisted genes **(Supplementary table 2).**

### 2.6 Statistical analysis

Statistical analysis was performed using GraphPad Prism software and median ± standard deviation (SD) was calculated. Differences between groups were assessed using an unpaired t-test (Mann-Whitney U test) and p value≤0.05 was considered statistically significant. GraphPad Prism version 9 was used as statistics software for validation data and for plotting representative graphs. Shapiro-Wilk test was used to check the distribution of data. Validation data was presented as median with interquartile range. P-value < 0.05 was considered as statistically significant.

## 3.0 Results

The demographic characteristics of the participants in the study were analysed. All the recruited OA subjects closely resembled the typical knee OA population. Among the participants with Grade 4 knee OA, we observed a notably higher median age and elevated pain scores assessed using the WOMAC (Western Ontario and McMaster Universities Osteoarthritis) scale (11). It was observed that individuals with advanced OA (Grade 4) exhibited a significant 62.15% increase in pain scores, while those with intermediate OA showed a 60.42% increase, both compared to individuals in the early stage (Grade 2) who had a pain score increase of 56.3% **(Table 1).**

### 3.1 Transcriptional profiles of differentially expressed genes

Microarray analysis was conducted using RNA isolated from synovial fluid samples from knee OA patients at different stages of the disease: grade 2 (n=3), grade 3 (n=3), and grade 4 (n=4). The RNA integrity numbers (RIN) for these RNA samples ranged from 7.2 to 9.6 **(Supplementary figure 2 and table 1)**. Each sample was treated as an independent data point, with no repeated measures, ensuring statistical independence.

The data normalization was performed using GeneSpring software, and ANOVA analysis was performed. Principal component analysis (PCA) reveals distribution of different grades of OA in different clusters, reflecting their unique transcriptomic profiles **(Figure 2).** The resulting hierarchal clustering heatmap visually depicted differential gene expression patterns across the OA samples, revealing three distinct clusters that correspond to the different OA grades **(Supplementary figure 3a).** The correlation analysis clearly demarcated the differences between the OA grades, highlighting significant distinctions in gene expression patterns **(Supplementary figure 3b).** Within each OA grade, patients showed positive correlations in gene expression, indicating similarity within groups. However, strong negative correlations were observed between different grades, underscoring significant intergrade differences.

**Figure 2.**
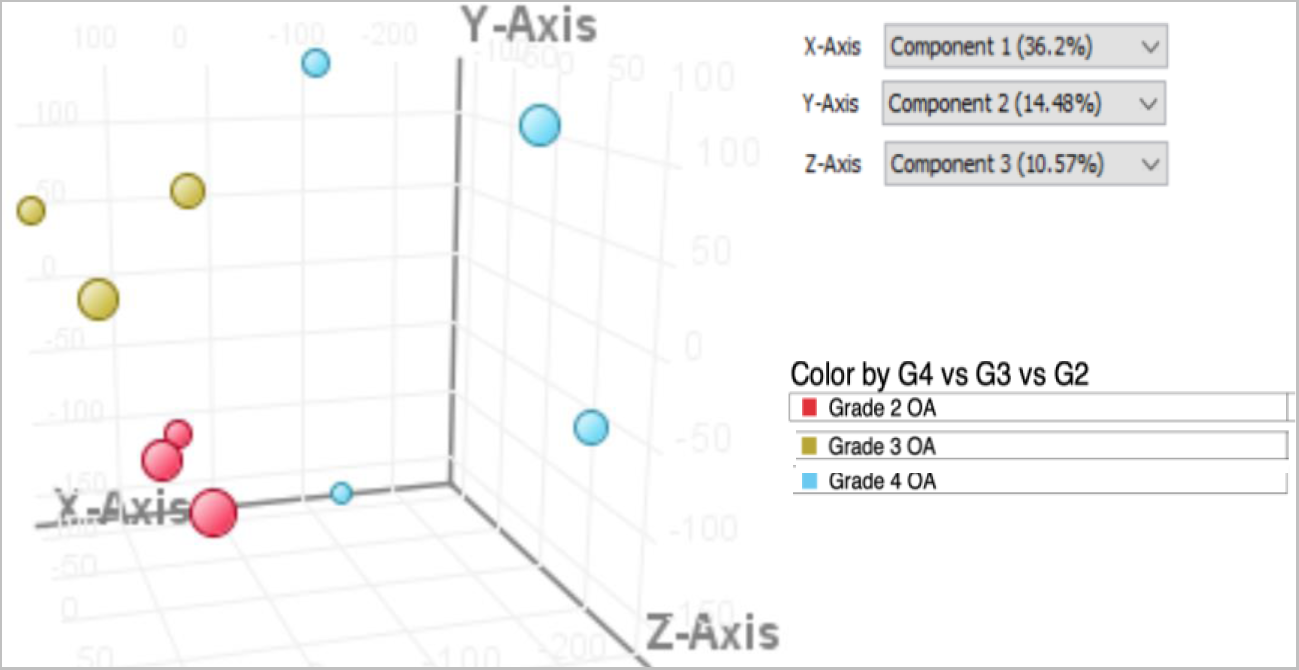
Principal component analysis plot of synovial fluid samples of different grades of osteoarthritis patients.

**Figure 3.**
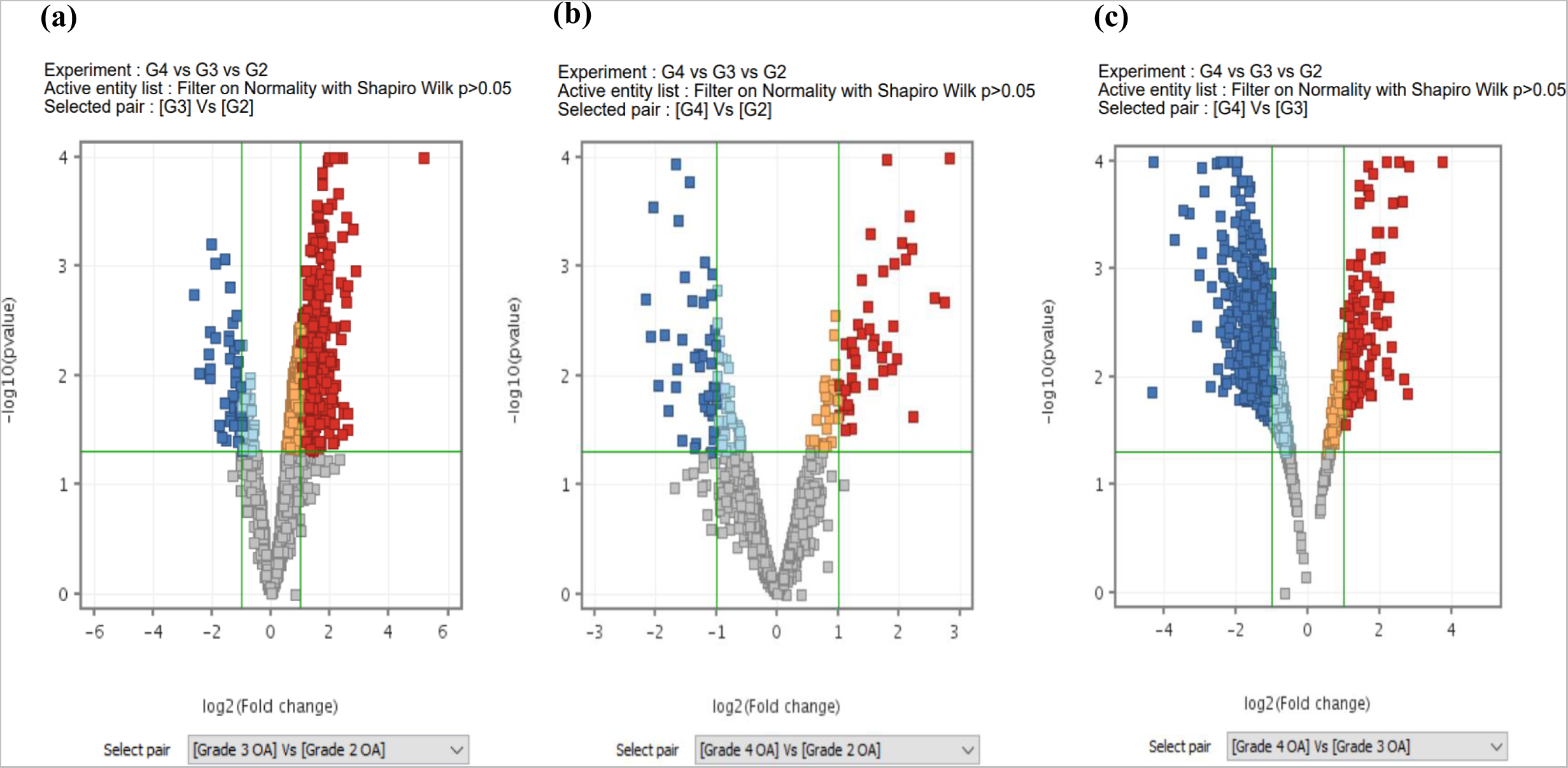
The volcano plot illustrates the distribution of differentially expressed genes in the intergrade analysis, specifically comparing (a) G3 *v/s* G2, (b) G4 *v/s* G3, and (c) G4 *v/s* G2, using moderated T-test. In this visual representation, genes which passed both cut-offs of fold change and p-value and are upregulated are depicted in red, genes which passed both cut-offs of fold change and p-value and are downregulated are shown in blue, genes in orange failed to pass fold change cut-off and are upregulated, genes in light blue failed to pass fold change cut-off and are downregulated and genes which failed to pass both cut-offs are represented in gray. (Fold change <1 or >-1; corrected *p-value* ≤0.05). G2-Grade 2, G3-Grade 3, G4-Grade 4.

To compare the progression of OA across different grades, an analysis of variance (*ANOVA*) test was performed in which late grades were compared with early grades so as to study the progression of the disease. This analysis focused on three comparison groups: Grade 3 vs Grade 2, Grade 4 vs Grade 2 and Grade 4 vs Grade 3. A total of 6500 genes showed differential expression between the grades (p-value ≤ 0.05 and fold change ≤ -1 or ≥ 1) **(Figure 4).**

**Figure 4.**
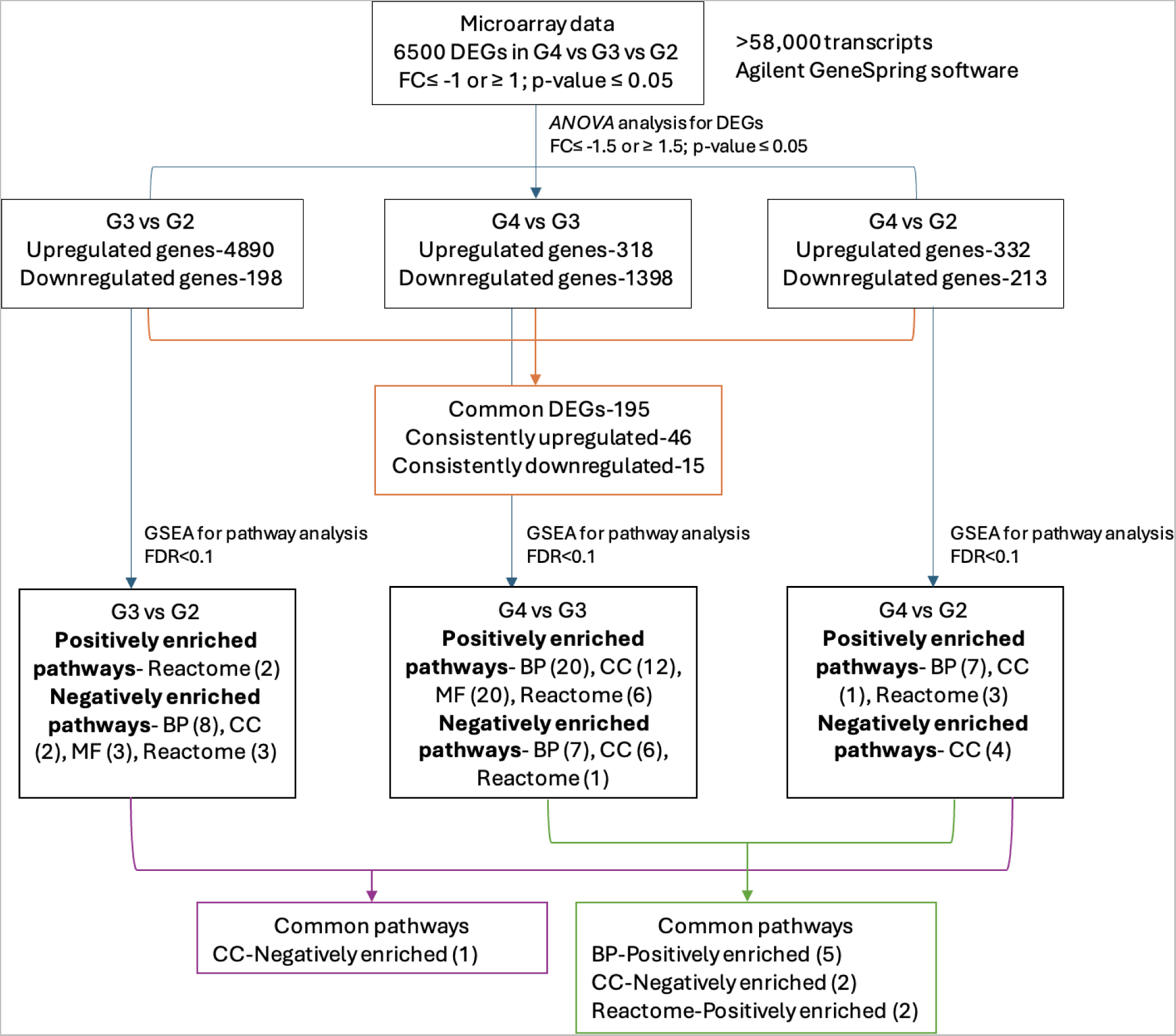
Flow chart of microarray data analysis. The agilent single color microarray platform was used having more than 58,000 transcripts and further data processing was done using agilent GeneSpring software followed by pathway analysis using GSEA database. Numbers in parentheses illustrates number of pathways enriched in each category using GSEA database. DEGs-Differentially expressed genes, ANOVA-Analysis of variance, BP-Biological pathway, CC-Cellular component, MF-Molecular function.

However, within each group screening with a stricter fold change criterion (≤ -1.5 or ≥ 1.5) revealed specific gene expression changes. In the Grade 4 vs Grade 2 comparison, 332 genes were upregulated and 213 genes were downregulated. In the Grade 4 vs Grade 3 comparison, 318 genes were upregulated, while 1398 genes were downregulated. For the Grade 3 vs Grade 2 comparison, 4890 genes were upregulated and 1198 genes were downregulated **(Figure 4 & Supplementary datasheet 1)**. These findings provide valuable insights into the genetic alterations associated with OA progression across different grades.

195 differentially expressed genes were found to be common in all three comparisons. However, 46 genes were found to be consistently upregulated and 15 were consistently downregulated with the progression of the disease (fold-change (≤ -1.5 or ≥ 1.5 and a p-value<0.05) **(Figure 4 & table 2).** These genes are important to study the progression of the disease from early to late stage.

**Table 2.**
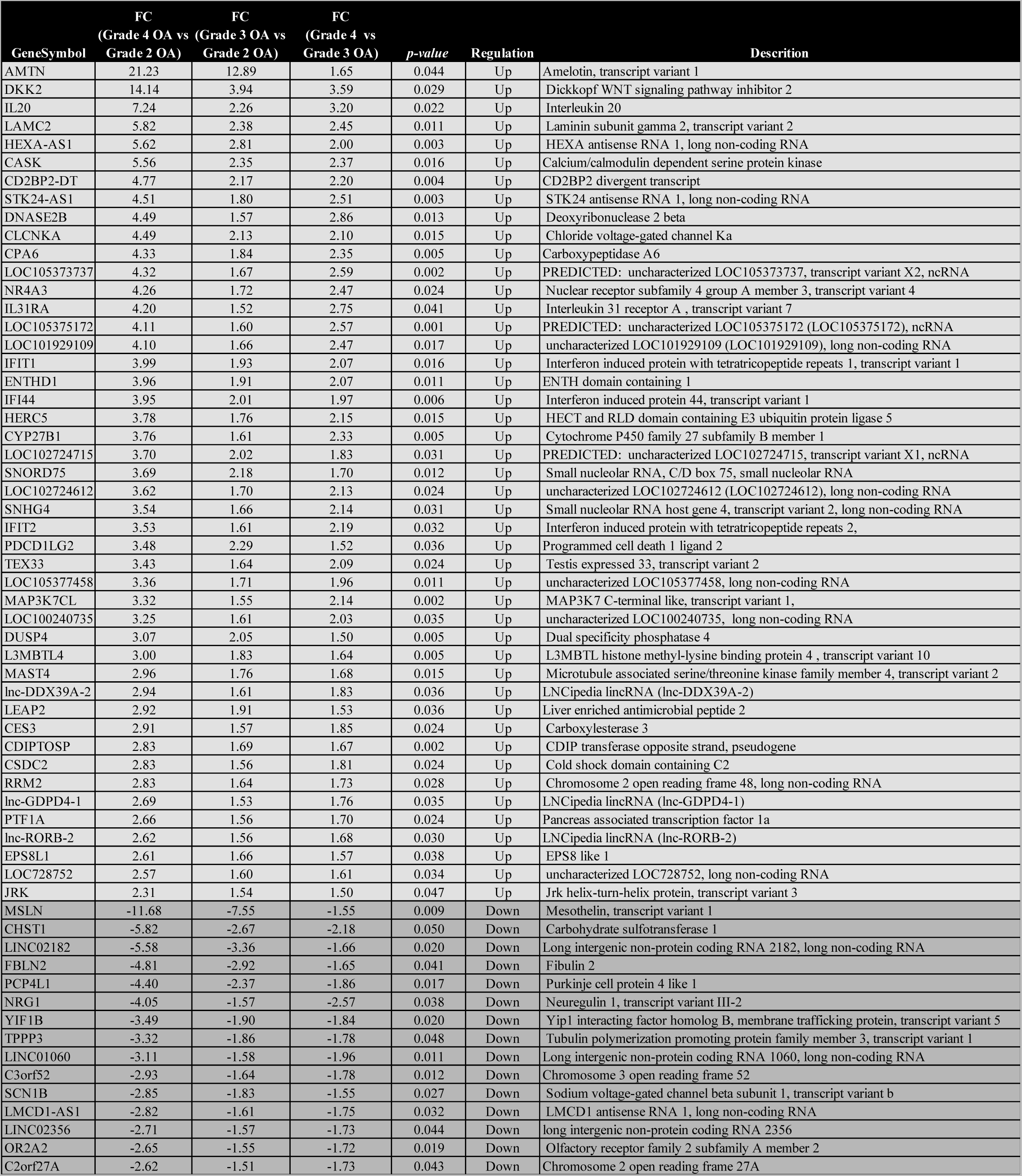
Common consistently upregulated and downregulated differentially expressed genes with the progression of the disease (FC≤ -1.5 or ≥ 1.5, p-value<0.05)

| GeneSymbol | FC<br>(Grade 4 OA vs<br>Grade 2 OA) | FC<br>(Grade 3 OA vs<br>Grade 2 OA) | FC<br>(Grade 4 vs<br>Grade 3 OA) | p-value | Regulation | Description |
| --- | --- | --- | --- | --- | --- | --- |
| AMTN | 21.23 | 12.89 | 1.65 | 0.044 | Up | Amelotin, transcript variant 1 |
| DKK2 | 14.14 | 3.94 | 3.59 | 0.029 | Up | Dickkopf WNT signaling pathway inhibitor 2 |
| IL20 | 7.24 | 2.26 | 3.20 | 0.022 | Up | Interleukin 20 |
| LAMC2 | 5.82 | 2.38 | 2.45 | 0.011 | Up | Laminin subunit gamma 2, transcript variant 2 |
| HEXA-AS1 | 5.62 | 2.81 | 2.00 | 0.003 | Up | HEXA antisense RNA 1, long non-coding RNA |
| CASK | 5.56 | 2.35 | 2.37 | 0.016 | Up | Calcium/calmodulin dependent serine protein kinase |
| CD2BP2-DT | 4.77 | 2.17 | 2.20 | 0.004 | Up | CD2BP2 divergent transcript |
| STK24-AS1 | 4.51 | 1.80 | 2.51 | 0.003 | Up | STK24 antisense RNA 1, long non-coding RNA |
| DNASE2B | 4.49 | 1.57 | 2.86 | 0.013 | Up | Deoxyribonuclease 2 beta |
| CLCNKA | 4.49 | 2.13 | 2.10 | 0.015 | Up | Chloride voltage-gated channel Ka |
| CPA6 | 4.33 | 1.84 | 2.35 | 0.005 | Up | Carboxypeptidase A6 |
| LOC105373737 | 4.32 | 1.67 | 2.59 | 0.002 | Up | PREDICTED: uncharacterized LOC105373737, transcript variant X2, ncRNA |
| NR4A3 | 4.26 | 1.72 | 2.47 | 0.024 | Up | Nuclear receptor subfamily 4 group A member 3, transcript variant 4 |
| IL31RA | 4.20 | 1.52 | 2.75 | 0.041 | Up | Interleukin 31 receptor A , transcript variant 7 |
| LOC105375172 | 4.11 | 1.60 | 2.57 | 0.001 | Up | PREDICTED: uncharacterized LOC105375172 (LOC105375172), ncRNA |
| LOC101929109 | 4.10 | 1.66 | 2.47 | 0.017 | Up | uncharacterized LOC101929109 (LOC101929109), long non-coding RNA |
| IFIT1 | 3.99 | 1.93 | 2.07 | 0.016 | Up | Interferon induced protein with tetratricopeptide repeats 1, transcript variant 1 |
| ENTHD1 | 3.96 | 1.91 | 2.07 | 0.011 | Up | ENTH domain containing 1 |
| IFI44 | 3.95 | 2.01 | 1.97 | 0.006 | Up | Interferon induced protein 44, transcript variant 1 |
| HERC5 | 3.78 | 1.76 | 2.15 | 0.015 | Up | HECT and RLD domain containing E3 ubiquitin protein ligase 5 |
| CYP27B1 | 3.76 | 1.61 | 2.33 | 0.005 | Up | Cytochrome P450 family 27 subfamily B member 1 |
| LOC102724715 | 3.70 | 2.02 | 1.83 | 0.031 | Up | PREDICTED: uncharacterized LOC102724715, transcript variant X1, ncRNA |
| SNORD75 | 3.69 | 2.18 | 1.70 | 0.012 | Up | Small nucleolar RNA, C/D box 75, small nucleolar RNA |
| LOC102724612 | 3.62 | 1.70 | 2.13 | 0.024 | Up | uncharacterized LOC102724612 (LOC102724612), long non-coding RNA |
| SNHG4 | 3.54 | 1.66 | 2.14 | 0.031 | Up | Small nucleolar RNA host gene 4, transcript variant 2, long non-coding RNA |
| IFIT2 | 3.53 | 1.61 | 2.19 | 0.032 | Up | Interferon induced protein with tetratricopeptide repeats 2, |
| PDCD1LG2 | 3.48 | 2.29 | 1.52 | 0.036 | Up | Programmed cell death 1 ligand 2 |
| TEX33 | 3.43 | 1.64 | 2.09 | 0.024 | Up | Testis expressed 33, transcript variant 2 |
| LOC105377458 | 3.36 | 1.71 | 1.96 | 0.011 | Up | uncharacterized LOC105377458, long non-coding RNA |
| MAP3K7CL | 3.32 | 1.55 | 2.14 | 0.002 | Up | MAP3K7 C-terminal like, transcript variant 1, |
| LOC100240735 | 3.25 | 1.61 | 2.03 | 0.035 | Up | uncharacterized LOC100240735, long non-coding RNA |
| DUSP4 | 3.07 | 2.05 | 1.50 | 0.005 | Up | Dual specificity phosphatase 4 |
| L3MBTL4 | 3.00 | 1.83 | 1.64 | 0.005 | Up | L3MBTL histone methyl-lysine binding protein 4 , transcript variant 10 |
| MAST4 | 2.96 | 1.76 | 1.68 | 0.015 | Up | Microtubule associated serine/threonine kinase family member 4, transcript variant 2 |
| lnc-DDX39A-2 | 2.94 | 1.61 | 1.83 | 0.036 | Up | LNCipedia lincRNA (lnc-DDX39A-2) |
| LEAP2 | 2.92 | 1.91 | 1.53 | 0.036 | Up | Liver enriched antimicrobial peptide 2 |
| CES3 | 2.91 | 1.57 | 1.85 | 0.024 | Up | Carboxylesterase 3 |
| CDIPTOSP | 2.83 | 1.69 | 1.67 | 0.002 | Up | CDIP transferase opposite strand, pseudogene |
| CSDC2 | 2.83 | 1.56 | 1.81 | 0.024 | Up | Cold shock domain containing C2 |
| RRM2 | 2.83 | 1.64 | 1.73 | 0.028 | Up | Chromosome 2 open reading frame 48, long non-coding RNA |
| lnc-GDPD4-1 | 2.69 | 1.53 | 1.76 | 0.035 | Up | LNCipedia lincRNA (lnc-GDPD4-1) |
| PTF1A | 2.66 | 1.56 | 1.70 | 0.024 | Up | Pancreas associated transcription factor 1a |
| lnc-RORB-2 | 2.62 | 1.56 | 1.68 | 0.030 | Up | LNCipedia lincRNA (lnc-RORB-2) |
| EPS8L1 | 2.61 | 1.66 | 1.57 | 0.038 | Up | EPS8 like 1 |
| LOC728752 | 2.57 | 1.60 | 1.61 | 0.034 | Up | uncharacterized LOC728752, long non-coding RNA |
| JRK | 2.31 | 1.54 | 1.50 | 0.047 | Up | Jrk helix-turn-helix protein, transcript variant 3 |
| MSLN | -11.68 | -7.55 | -1.55 | 0.009 | Down | Mesothelin, transcript variant 1 |
| CHST1 | -5.82 | -2.67 | -2.18 | 0.050 | Down | Carbohydrate sulfotransferase 1 |
| LINC02182 | -5.58 | -3.36 | -1.66 | 0.020 | Down | Long intergenic non-protein coding RNA 2182, long non-coding RNA |
| FBLN2 | -4.81 | -2.92 | -1.65 | 0.041 | Down | Fibulin 2 |
| PCP4L1 | -4.40 | -2.37 | -1.86 | 0.017 | Down | Purkinje cell protein 4 like 1 |
| NRG1 | -4.05 | -1.57 | -2.57 | 0.038 | Down | Neuregulin 1, transcript variant III-2 |
| YIF1B | -3.49 | -1.90 | -1.84 | 0.020 | Down | Yip1 interacting factor homolog B, membrane trafficking protein, transcript variant 5 |
| TPPP3 | -3.32 | -1.86 | -1.78 | 0.048 | Down | Tubulin polymerization promoting protein family member 3, transcript variant 1 |
| LINC01060 | -3.11 | -1.58 | -1.96 | 0.011 | Down | Long intergenic non-protein coding RNA 1060, long non-coding RNA |
| C3orf52 | -2.93 | -1.64 | -1.78 | 0.012 | Down | Chromosome 3 open reading frame 52 |
| SCN1B | -2.85 | -1.83 | -1.55 | 0.027 | Down | Sodium voltage-gated channel beta subunit 1, transcript variant b |
| LMCD1-AS1 | -2.82 | -1.61 | -1.75 | 0.032 | Down | LMCD1 antisense RNA 1, long non-coding RNA |
| LINC02356 | -2.71 | -1.57 | -1.73 | 0.044 | Down | long intergenic non-protein coding RNA 2356 |
| OR2A2 | -2.65 | -1.55 | -1.72 | 0.019 | Down | Olfactory receptor family 2 subfamily A member 2 |
| C2orf27A | -2.62 | -1.51 | -1.73 | 0.043 | Down | Chromosome 2 open reading frame 27A |

Among the consistently upregulated genes, *AMTN* (Amelotin) and *DKK2* (Dickkopf WNT Signaling Pathway Inhibitor 2) are the two top most upregulated genes, whereas, *MSLN* (Mesothelin) is the top downregulated gene with the progression of the disease having foldchange cut-off more than 10 folds in late stages of the disease. *AMTN,* a gene involved in hydroxyapatite mineralization during enamel maturation, is typically released by ameloblast cells (12). In the current study, *AMTN* emerged as the most upregulated gene in synovial fluid samples from OA patients, with its expression correlating with the severity of the disease, particularly in advanced stages. Specifically, *AMTN* expression consistently increased in late-stage OA, with grades 3 and 4 showing a 12.89 and 21.23 fold upregulation, respectively, compared to early-stage grade 2. Another noteworthy gene, *DKK2* has a significant role in terminal osteoblast differentiation into mineralized matrices (13). *DKK2* is also dysregulated in OA, with its expression significantly increasing in the later stages of the disease. In late-stage OA grade 4, *DKK2* expression is 14.14-fold higher compared to early-stage grade 2, indicating its involvement in the progression and severity of OA.

Mesothelin (*MSLN*) is the top most downregulated gene with its fold change as low as -11.68 and -7.55 in late grade 4 and grade 3 OA when compared to early grade 2 OA. *Mesothelin* is reported to be involved in bone remodelling and suggested as a potential therapeutic target for the treatment of bone disorders (14).

### 3.2 Analysis of Gene Ontology enrichment

Pathway analysis was conducted to study the dysregulated pathways and their gene counterparts with the progression of the disease. Specifically, comparison of late grades was done with early grades to discern patterns of pathway alterations as the disease advances. This comparison allowed to track how key pathways evolve over the course of disease progression, providing insights into the biological, cellular and molecular changes underlying the severity of OA. From GSEA database, the list of positively and negatively enriched pathways with their association with the disease were analyzed based on the gene ontology (GO) biological pathways, cellular components, and molecular functions. Furthermore, DEGs were also analyzed using reactome pathways **(Figure 4).**

The comparative analysis of OA progression from grade 2 to grade 3 reveals significant molecular changes. Notably, the upregulation of reactome pathways such as neutrophil degranulation and developmental biology, indicates an enhanced inflammatory response and changes in tissue growth and repair mechanisms **(Figure 5 & Supplementary table 3)**. Conversely, several pathways are downregulated, including those involved in autophagy, programmed cell death, and inflammatory response regulation, RNA processing, transcriptional regulation, and cellular components like the golgi apparatus and nucleolus points to disruptions in gene expression and protein processing. Overall, these molecular changes underscore the complex interplay between inflammation, impaired cellular mechanisms, and tissue degradation in OA progression **(Figure 5 & Supplementary table 3).**

**Figure 5.**
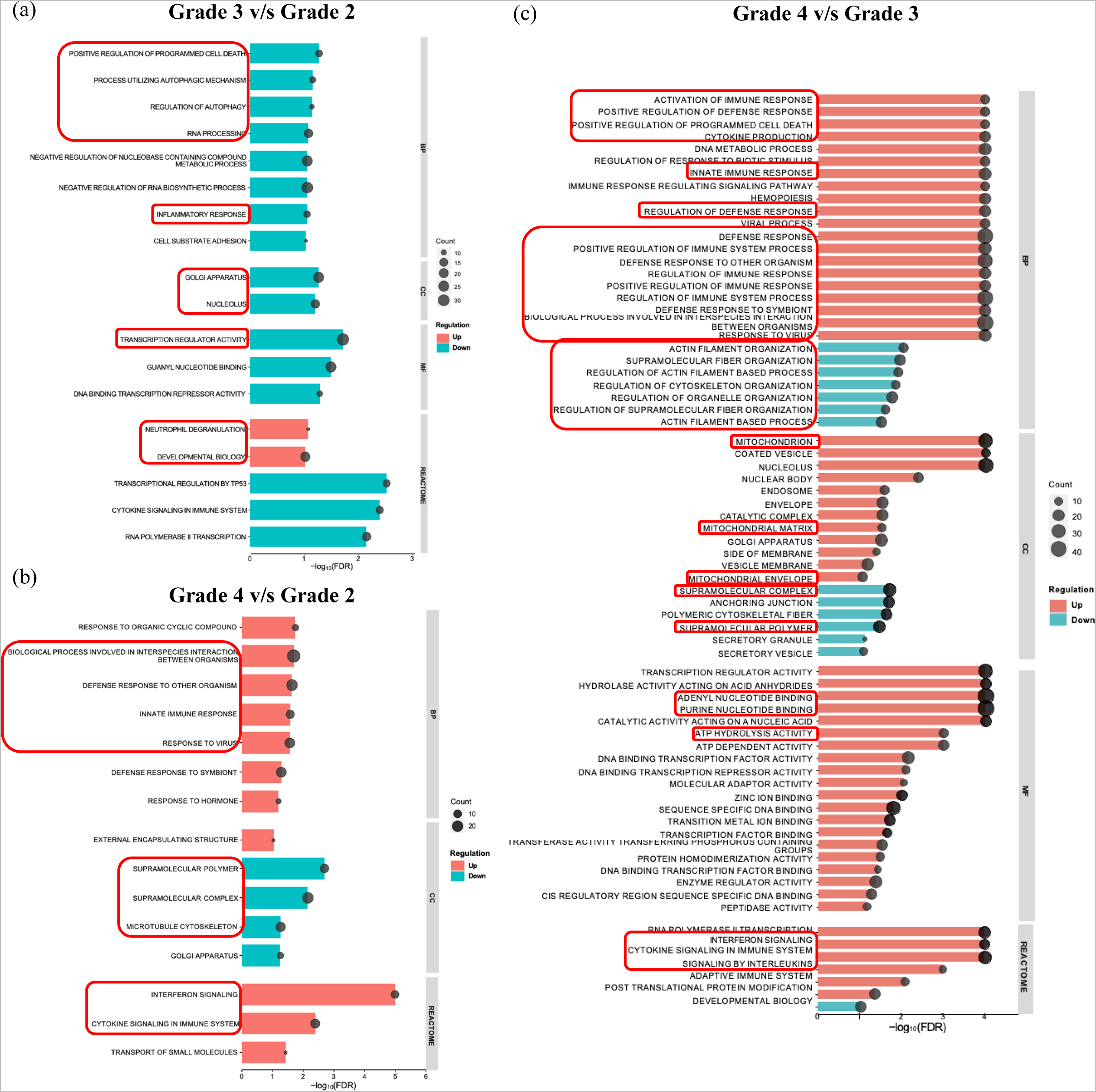
GO bar plot of biological, cellular component, molecular function and reactome of differentially expressed upregulated and downregulated pathways between (a) Grade 3 *v/s* Grade 2 (b) Grade 4 *v/s* Grade 2 (c) Grade 4 *v/s* Grade 3 osteoarthritis group with the GSEA database. (*p-value*≤0.05, FDR≤0.1). The bar plots has been made using SR plot database. Encircled pathways have been discussed in the text. Highlighted pathways are discussed in the text.

The comparative analysis of biological pathways between grade 4 and grade 2 OA reveals significant changes, reflecting disease progression. In advanced OA, there is a marked upregulation in immune and inflammatory pathways. The defense response to symbiont pathway, indicates an elevated immune response against microbial invaders. The biological process involved in interspecies interaction suggests extensive molecular interactions, possibly with the microbiome. Similarly, the response to virus pathway and interferon signaling pathway highlight an increased antiviral response and robust activation of interferon pathways, critical for immune regulation. Additionally, the enrichment of reactome pathways like cytokine signaling in immune system pathway reflects enhanced cytokine activity, contributing to chronic inflammation observed in advanced OA **(Figure 5 & Supplementary table 4)**. Conversely, pathways associated with cellular structure and integrity are downregulated in advanced OA reflecting compromised structural support and disruptions in protein processing, essential for maintaining cell shape and function. These molecular changes underscore the complex interplay between inflammation and cellular degradation in the progression of OA, highlighting the dual impact of increased immune activity and structural decline in advanced stages of the disease **(Figure 5 & Supplementary table 4).**

In the comparative analysis of grade 4 vs grade 3 OA several positively enriched biological pathways such as defense response to other organism, innate immune response, response to virus and biological process involved in interspecies interaction points to an enhanced inflammatory response to microbial invasion and viral infections in advanced OA. Other notable pathways include regulation of immune system process, positive regulation of immune response and defense response to other organism, thus indicating increased immune response in the advanced stage of disease. Additionally, reactome pathways like interferon and cytokine production suggests high-grade synovial inflammation or synovitis in advanced stages of the disease. There is significant positively enriched pathways in cellular components like mitochondria, including their matrix and envelope seen in end stage OA suggesting mitochondrial dysfunctioning **(Figure 5 & Supplementary table 5)**. Alongwith, the molecular pathways like ATP hydrolysis, adenyl, and purine nucleotide binding pathways are also upregulated indicating a possible disruption of mitochondrial activity leading to upregulation of several inflammation related pathways. Similar to grade 4 vs 2 analysis, cellular components like supramolecular polymer and complex were downregulated. As many of the supramolecular compounds like collagen and proteoglycan gets degraded as the disease progresses and gets completely degraded in advanced OA indicating a decrease in these supramolecular complexes. Downregulation of actin and cytoskeleton components implies alterations in cellular structure and mechanical support within joint tissues. Notably, the downregulation of tubulin, a key component of microtubules involved in cell division and intracellular transport, indicates disruptions in these essential cellular processes in late stage OA **(Figure 5 & Supplementary table 5).**

Overall, OA progression is marked by the upregulation of inflammation-related pathways and the downregulation of cellular structural pathways. This highlights the potential for targeting inflammation and cytoskeletal organization in therapeutic interventions for OA.

### 3.3 Common pathways involved across the disease progression

The progression of OA from Grade 2 to Grade 3, Grade 2 to Grade 4, and Grade 3 to Grade 4 involves disruptions in cellular components and downregulation of pathways related to supramolecular complexes and polymers. As OA progresses, supramolecular compounds such as collagen and proteoglycan degrade, nearly disappearing in advanced stages, indicating a decrease in these complexes. Early-stage disruptions in cellular components, like the Golgi apparatus, suggest protein misfolding. This misfolding triggers an inflammatory response as OA advances, supported by the enrichment of immune-related pathways in Grade 4, the disease’s end stage.

Moreover, defense response pathways related to symbionts, viruses, and interspecies interactions highlight the presence of external microbes in later OA stages. The overall elevation of immune-regulated pathways and the decline in pathways related to cellular structure and integrity in advanced OA suggest these pathways as potential targets for therapeutic interventions **(Figure 4 & table 3).**

**Table 3.** List of common GSEA pathways in intergrade comparison between different grades of osteoarthritis.

| Pathway | GO term | Enrichment | Grade 3 vs Grade 2 | Grade 4 vs Grade 2 | Grade 4 vs Grade 3 |
| --- | --- | --- | --- | --- | --- |
| Defense response to symbiont | BP | Positive | - | + | + |
| Response to Virus | BP | Positive | - | + | + |
| Biological process involved in interspecies interaction between organisms | BP | Positive | - | + | + |
| Defense response to other organism | BP | Positive | - | + | + |
| Innate immune response | BP | Positive | - | + | + |
| Cytokine signaling in immune system | Reactome | Positive | - | + | + |
| Interferon signaling | Reactome | Positive | - | + | + |
| Supramolecular polymer | CC | Negative | - | + | + |
| Supramolecular complex | CC | Negative | - | + | + |
| Golgi apparatus | CC | Negative | + | + | - |

+Enriched, -Not enriched, GO-Gene ontology, BP-Biological pathway, CC-Cellular component

### 3.4 STRING analysis

To study the progression of the disease, comparing Grade 4 to Grade 2 is ideal as it contrasts late and early OA stages, therefore gene signatures of the affected pathways from grade 4 v/s grade 2 comparison were analyzed. Interactome studies were conducted to examine the protein-protein interaction networks of these gene counterparts with a PPI enrichment *p-value*<1.0e-16 and a high interaction score confidence of 0.7. This analysis resulted in the formation of five clusters, with the majority of genes located in cluster 1 **(Figure 6a).** Further pathways from cluster 1 were analyzed. The most densely connected pathway with the highest strength belong to defense response to virus in biological pathways with a highly significant FDR of 6.34e-20. Molecular function pathways reveal double stranded RNA binding as one of the enriched pathways with a FDR of 4.55e-06. Reactome pathways with highest FDR of 3.17e-14 belong to interferon signaling **(Figure 6b).** These results further supported the data from GSEA database indicating that increased inflammation in late stages of OA is a key manifestation of the disease.

**Figure 6.**
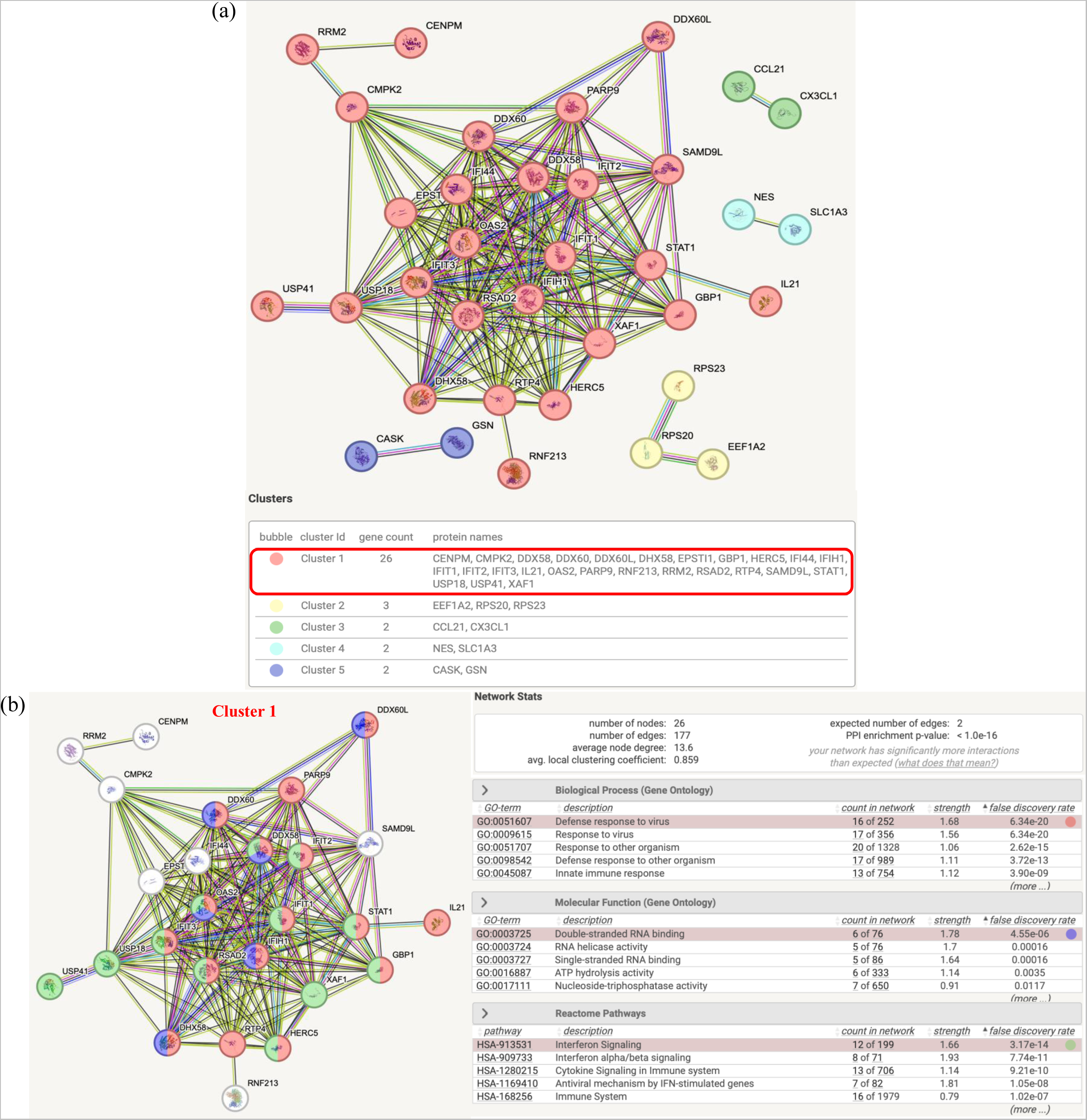
(a) Protein-protein interaction networks of the differentially expressed genes from Grade 4 *v/s* Grade 2 comparison using STRING database; (b) Network of genes from cluster 1 and its related pathways. Disconnected nodes were removed.

### 3.5 qRT-PCR validation

For validation of microarray data, genes which were found to be consistently upregulated with the progression of the disease were selected. *AMTN and DKK2* being the top two upregulated genes found to increase with the progression of the disease and are present in all three intergrade comparisons (grade 3 vs grade 2, grade 4 vs grade 2 and grade 4 vs grade 3).

Furthermore, to validate the findings of GSEA pathway analysis and interactome studies, genes belonging to the consistently upregulated pathways like defense response, cytokine and interferon signaling were also selected. *IFIT2, IFIT3 and STAT1* were selected due to their coexistence in all three pathways in end stage vs early stage of OA.

The study successfully validated the top upregulated genes, *AMTN* and *DKK2*, whose elevated expression was consistent with the microarray data. Additionally, the expression of genes *IFIT2, IFIT3,* and *STAT1* was also validated, demonstrating an upregulation in Grade 4 OA synovial fluid samples compared to Grade 2 samples **(Figure 7).**

**Figure 7.**
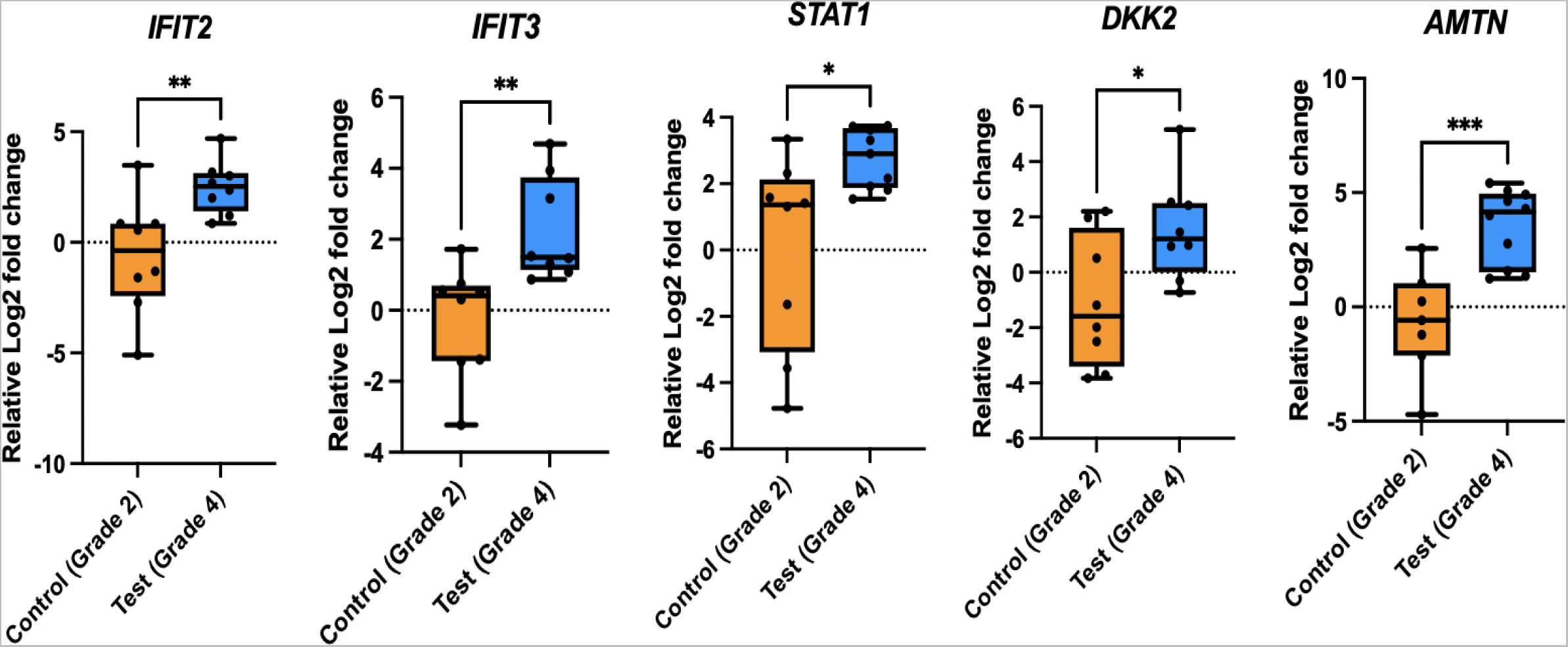
Relative expression analysis of (A) *IFIT2*, *IFIT3*, *STAT1, DKK2 and AMTN* demonstrating relative Log2 fold change expression in synovial fluid samples of Grade 4 v/s Grade 2 osteoarthritis patients. **p<0.01, ***p<0.001, ****p<0.0001. G2 OA(n=8) and G4 OA (n=10)

As *AMTN* is the common top most upregulated gene among all the three intergrade comparisons, protein quantification of *AMTN* was also done to check for its protein expression in synovial fluid samples of grade 2 and grade 4 OA patients. The results reveal an increase in protein levels of amelotin protein in end stage OA as compared to early stage indicating it as a protein to look forward for therapeutic interventions **(Supplementary figure 4).**

## 4.0 Discussion

Current study aimed to investigate the transcriptional and molecular changes associated with OA progression, focusing on the differential gene expression and pathway enrichment across various OA grades so as to study the progression of the disease from early to late stages.

Microarray analysis of synovial fluid samples revealed three distinct clusters corresponding to OA grades, with principal component analysis confirming these differences. A significant number of genes showed differential expression between the grades. The consistent upregulation of *AMTN* and *DKK2,* and the downregulation of *MSLN* with the progression of the disease across the intergrade analysis, was notable. *AMTN* is expressed by ameloblast cells in maturation stage of enamel formation. Its expression has been restricted to ameloblasts cells. Rajapakse et al, has revealed that the formation and deposition of hydroxyapatite mineral deposits in retinal pigment epithelial cells are associated with the progression of age-related macular degeneration.(15). Moreover, there is formation of pathological hydroxyapatite crystals in OA as well (16, 17). The formation of hydroxyapatite crystals in late-stage OA triggers an inflammatory cascade (17,18). Therefore, the increased *AMTN* expression with disease progression could contribute to the pathological formation of hydroxyapatite crystals and subsequent inflammation in OA. Although, previously elevated *AMTN* levels in the cartilage and synovium of OA patients have been documented, this study is the first report for the demonstration of upregulated *AMTN* gene expression (Figure 7) as well as protein **(Supplementary figure 4)** in synovial fluid of OA patients with progression of disease. Additionally, no studies had validated *AMTN* protein levels. *DKK2,* a Wnt signaling antagonist, was also upregulated (19). There are reports supporting the presence of *DKK2* proteomic signatures in synovial fluid of OA patients (20). *DKK2* is a potent antagonist for canonical Wnt signaling. The Wnt signaling pathway is crucial for osteogenesis and the regulation of terminal differentiation in osteoblasts.(21). In OA, Wnt signaling is disrupted. It is increased in OA chondrocytes leading to chondrocyte hypertrophy and mineralization whereas in sub-chondral bone there is decreased Wnt signaling and therefore osteopenic phenotype. This dichotomy highlights the complex nature of OA, where soft tissues undergo pathological mineralization, while subchondral bone experiences bone resorption due to disrupted signaling pathways. Synovial fluid, acting as a reservoir for these secreted components, reflects the pathological changes occurring within the joint, providing critical insights into the molecular mechanisms driving OA progression.

Conversely, *MSLN,* which is essential for osteoblast differentiation and function, was downregulated, indicating disrupted bone remodelling in OA (14). Studies have revealed that mesothelin knockout mice has osteoporotic phenotype. Subchondral bone in OA is known to accelerate osteoclastogenesis. *MSLN* deficiency leads to attenuated osteoblast differentiation and function which furthermore supports the decreased mineralization in subchondral bone (22).

The aspect of mineralization in OA still remains underexplored. These findings suggest a niche for therapeutic interventions targeting mineralization as a key manifestation of the disease, offering a novel avenue for treatment.

Further, pathway enrichment analysis highlighted significant changes with OA progression. As OA progresses from grade 2 to grade 3, neutrophil degranulation the topmost upregulated pathway involves the release of granular contents from neutrophils, which gets infiltrated to the synovial fluid and joint space in early OA due to joint tissue damage. Upon activation, neutrophils release enzymes and reactive oxygen species that contribute to an inflammatory environment, degrading cartilage and other joint structures, thereby amplifying tissue damage (23).

In end stage OA, i.e., grade 4 OA, pathways like defense response to other organism, innate immune response, and response to virus points to an enhanced inflammatory response to microbial invasion and viral infections as compared to grade 3 and grade 2 OA. Various studies support the presence of angiogenesis in end stage OA (24). Microbial components, like bacterial endotoxins, can infiltrate joint tissues through bloodstream infections or local invasions, triggering immune responses that increase inflammation and joint damage. The body’s defense involves activating immune cells and releasing inflammatory mediators, contributing to the overall inflammatory environment in OA-affected joints. In OA, damage associated molecular patterns (DAMPs) from injured cartilage and other joint tissues are released into the synovial fluid (25). These DAMPs are recognized by pattern recognition receptors (PRRs) on synovial macrophages and other infiltrating immune cells, leading to cell activation and the production of pro-inflammatory cytokines (e.g., IL-1β, TNF-α, IL-6). This synovial inflammation aggravates joint damage and pain (26, 27). A notable trend was observed in the regulation of inflammatory response and cytokine signaling pathways. These pathways were initially downregulated in grade 3, likely as a protective mechanism to prevent excessive inflammation. However, in grade 4 OA, they were upregulated, indicating severe inflammation. The enrichment of interferon and cytokine signaling pathways in advanced stages suggests high-grade synovial inflammation or synovitis.

Furthermore, pathways such as programmed cell death, autophagy, and golgi function are downregulated in grade 3 OA. These pathways are interconnected, as autophagy is a form of programmed cell death. Uncontrolled programmed cell death poses a risk for secondary OA, while autophagy can mitigate OA progression and protect cartilage by removing damaged cellular components such as misfolded proteins and senescent organelles (28, 29). Autophagy process is initiated with the involvement of endoplasmic reticulum and the trans-Golgi network, helping in the removal of protein aggregates and damaged organelles by fusing them with lysosomes. The downregulation of autophagy pathways hampers the removal of these damaged proteins, cells, and organelles, leading to the accumulation of DAMPs (30, 31). When secreted into synovial fluid, DAMPs contribute to inflammation (25). Therefore, the dysregulation of these cellular processes exacerbates the inflammatory and degenerative aspects of OA, underscoring the significant role of inflammation in the progression and end-stage of OA.

Apart from the involvement of biological pathways related to inflammation, certain synovial fluid signature genes in OA were found to be primarily localized in mitochondria. Mitochondrial dysfunction in OA disrupts autophagy, leading to reactive oxygen species (ROS) production in chondrocytes. Impaired autophagy results in damaged mitochondria accumulation and activation of inflammatory pathways, contributing to OA’s pathogenesis and progression. Additionally, mitochondrial dysfunction compromises ATP production (32). Mitochondrial stress and chondrocyte senescence result in the release of mitochondrial double-stranded RNAs (mt-dsRNAs) into the cytosol. These mt-dsRNAs can trigger the innate immune system, thereby activating interferon signaling (33). Similar phenotypes have been seen in the prevailing data as well. There is upregulation of cellular components like mitochondria, including their matrix and envelope seen in end stage OA. Additionally, pathways like double stranded RNA binding consisting of key genes like *DDX60L, DDX60, DDX58, DHX58, IFIH1, RSAD2 and IFIT1* belong to a family of RNA sentinels (34). Alongwith, the molecular pathways like ATP hydrolysis, adenyl and purine nucleotide binding pathways are also upregulated indicating a possible disruption of mitochondrial activity leading to upregulation of several inflammation related pathways. (35)

In contrast cellular components like supramolecular polymer and complex were downregulated in grade 4 OA as compared to grade 2 and grade 3. As many of the supramolecular compounds like collagen and proteoglycan gets degraded as the disease progresses and gets completely degraded in advanced OA indicating a decrease in these supramolecular complexes (36). OA also causes changes in the cellular cytoskeleton, including the downregulation of actin and microtubule components. This implies alterations in cellular structure and mechanical support within joint tissues. Notably, the downregulation of tubulin, a key component of microtubules involved in cell division and intracellular transport, indicates disruptions in these essential cellular processes in OA (37).

The findings of our study shed light on the critical roles played by mineralization-related genes and inflammation-related pathways in the progression of OA from early to advance stages. Understanding how these genes and pathways are involved at different stages of the disease could pave the way for significant breakthroughs in OA research. By decoding the underlying mechanisms, we can identify potential targets for therapeutic interventions aimed at effectively managing bone remodeling and inflammatory processes in OA. The current study holds immense promise for improving patient care. By delving deeper into these pathways, we can develop more targeted and personalized treatment approaches. Ultimately, this could lead to better outcomes for individuals suffering from OA, enhancing their quality of life and offering hope for a brighter future.

## Contributions

**Rinkle Sharma:** Conception and design, analysis and interpretation of data, drafting of the article, statistical expertise, collection and assembly of data. **Diksha Rana:** Collection and assembly of data. **Rahul Kumar:** Statistical expertise. **Sakshi Narula**-Conception and design. **Alpa Chaudhary:** Conception and design. **Bhavneet Kaur:** Collection and assembly of data. **Khushpreet Kaur:** Conception and design. **Mandeep Singh Dhillon:** Provision of study materials or patients. **Devendra K Chauhan:** Provision of study materials or patients. **Uttam Chand Saini:** Provision of study materials or patients. **Sadhna Sharma:** Administrative, technical, or logistic support. **Jyotdeep Kaur:** Administrative, technical, or logistic support. **Indu Verma:** Conception and design, analysis and interpretation of data, critical revision of the article for important intellectual content, final approval of the article, obtaining of funding, administrative, technical, or logistic support.

## Supporting information

Supplementary data sheet 1

## Acknowledgments

This work was financially supported by Indian Council of Medical Research (ICMR), project No. 5/4/-5/3/TF/Ortho/2017-NCD-1.

## Data sharing statement

The data that support the findings of this study are publicly available in gene expression omnibus (GEO) dataset vide accession no. GSE262903.

