## Supplementary document 1 for "Synovial fluid transcriptome dynamics in osteoarthritis progression: Implications in pathogenesis"

**Supplementary material**

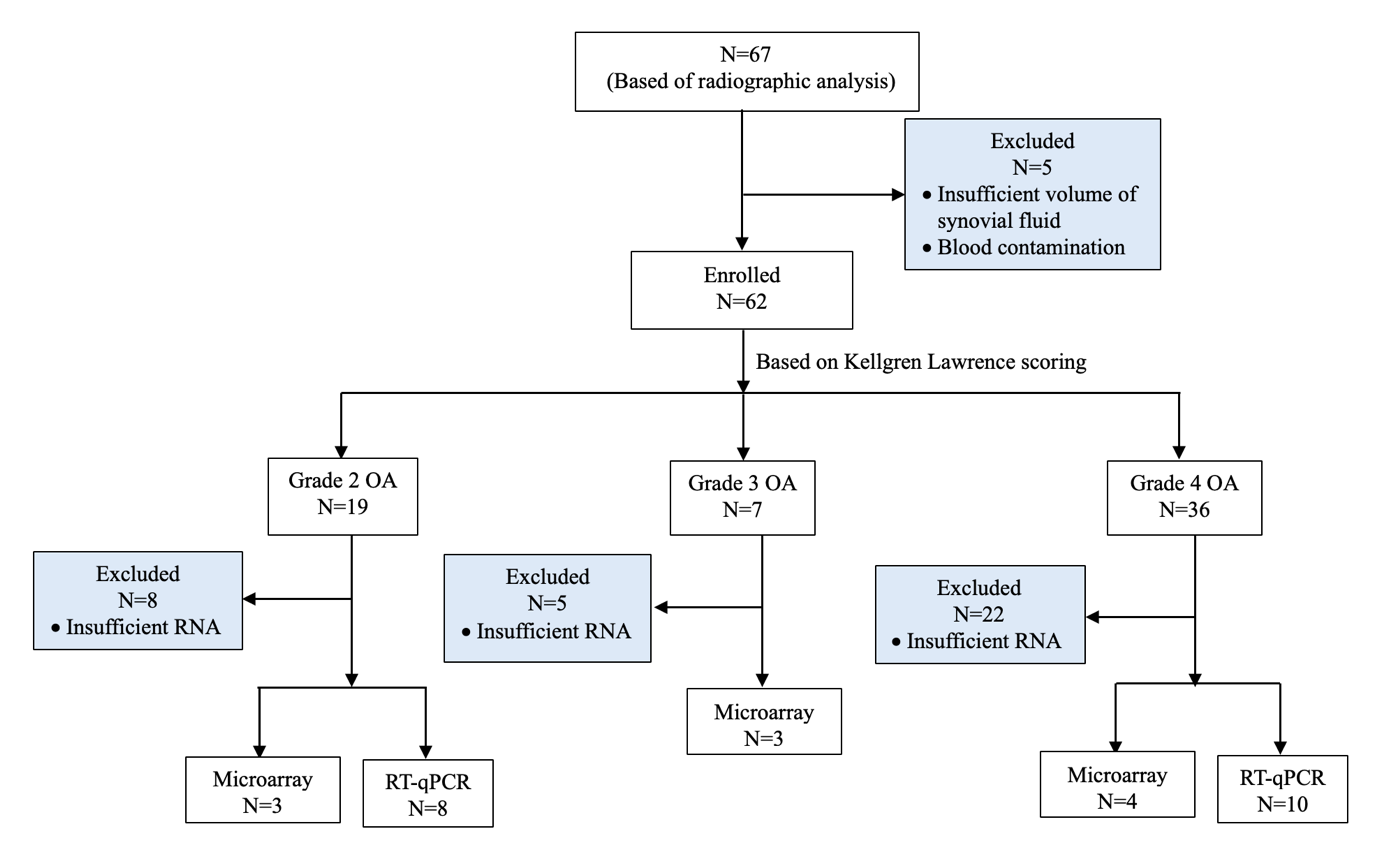

**Figure 1.** Consort diagram for flow of OA patients undergoing PRP/TKR treatment.

PRP-Platelet rich plasma, TKR-Total knee arthroplasty

| **Samples** | **RIN** |
| --- | --- |
| OA GRADE 2 (1) | 7.2 |
| OA GRADE 2 (2) | 8.1 |
| OA GRADE 2 (3) | 8.3 |
| OA GRADE 3 (1) | 8.8 |
| OA GRADE 3 (2) | 7.8 |
| OA GRADE 3 (3) | 8.9 |
| OA GRADE 4 (1) | 9.1 |
| OA GRADE 4 (2) | 8.7 |
| OA GRADE 4 (3) | 8.5 |
| OA GRADE 4 (4) | 9.6 |

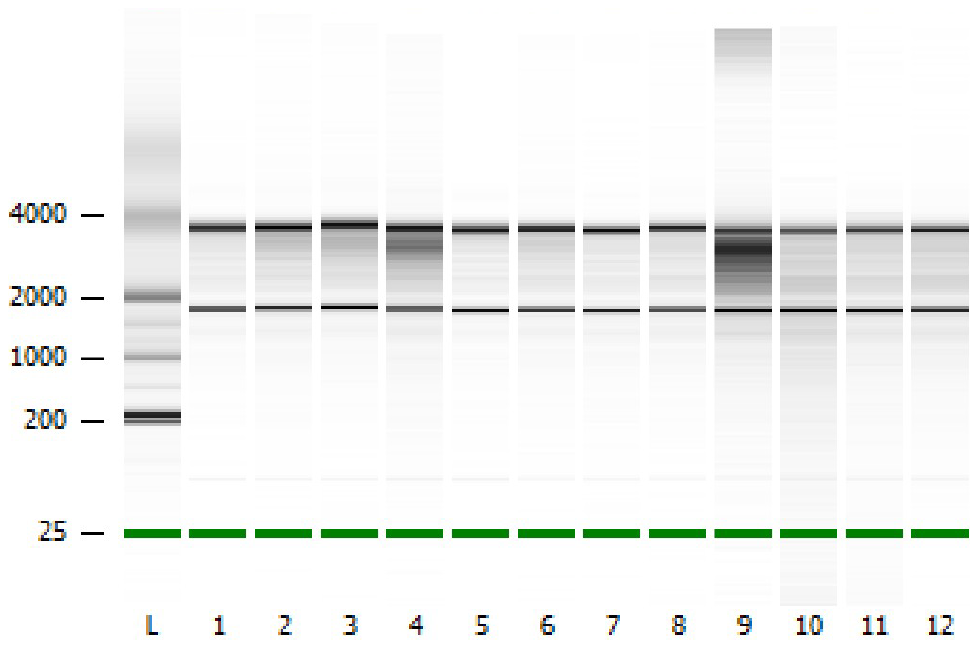

**Figure 2 and Table 1**. Representative image and RIN values different grades of synovial fluid samples of osteoarthritis patients assessed on bioanalyzer.

RIN-RNA integrity number

**Table 2.** Primer sequences of genes shortlisted from microarray and validated using qRT-PCR.

| Gene symbol | Forward primer sequence (5’-3’) | Forward primer sequence (5’-3’) | Annealing temperature |
| --- | --- | --- | --- |
| *GAPDH* | TCCCTGAGCTGAACGGGAAG | GGAGGAGTGGGTGTCGCTGT | 61.4 |
| *IFIT2* | ACAAGGCCATCCACCACTTT | TCCAGACTCCAAACCCCTCT | 60.6 |
| *IFIT3* | CCCTGGAATGCTTACGGCAA | CAGGCGTAGTTTCCCCAAGT | 60.6 |
| *STAT1* | GTGAAGTTGAGAGATGTGAATGAG | GATCACCACAACGGGCAGA | 52.8 |
| *DKK2* | CCCAGTACCCGCTGCAATAA | GTGCCGAGTACCATCCAGAG | 61.6 |
| *AMTN* | GTTGAATGTACAACAGCAACTGCAC | TTCCATCCTGGACATCTGGATTAG | 61.4 |

.

(a)

(b)

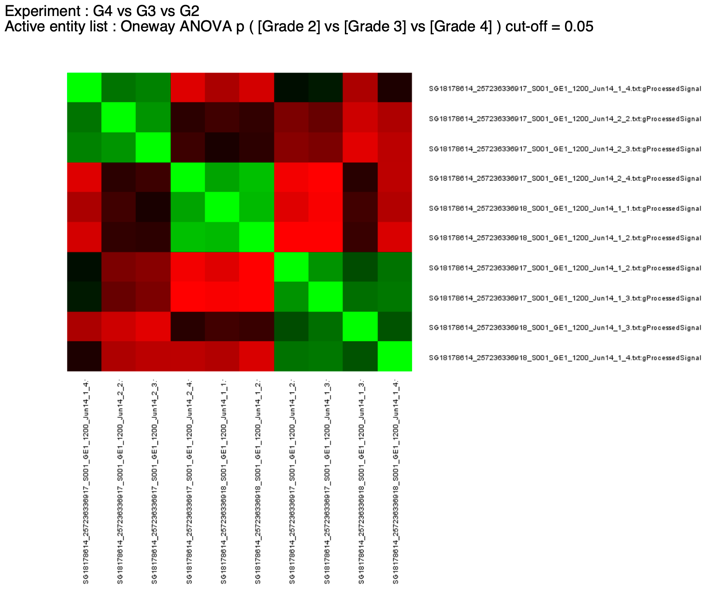

**Grade 3**

**Grade 2**

**Grade 4**

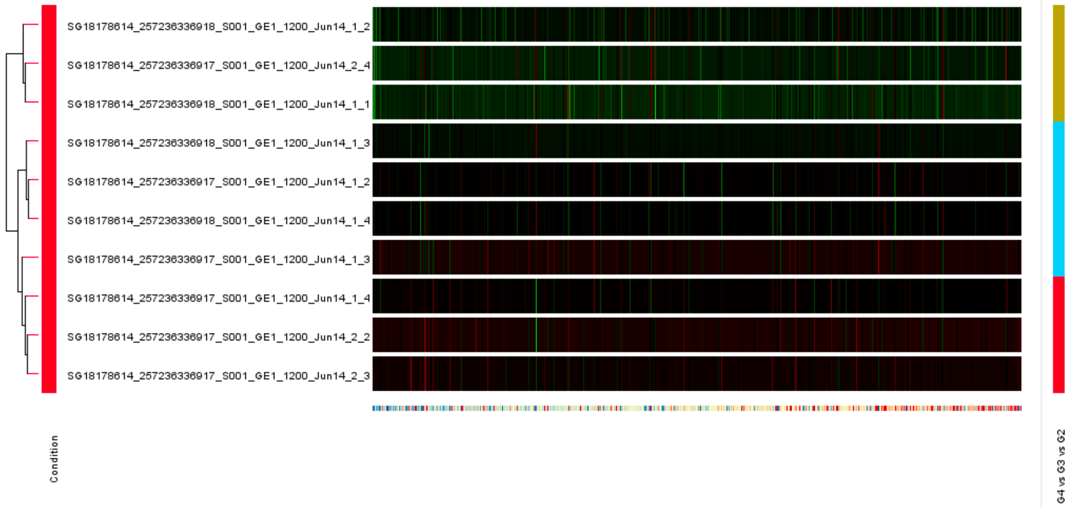

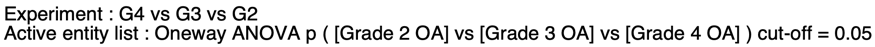

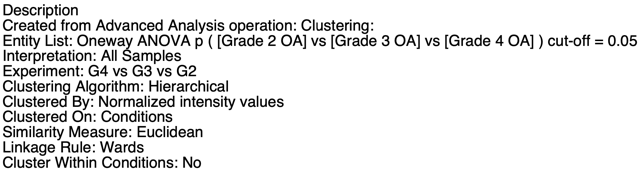

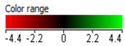

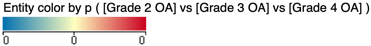

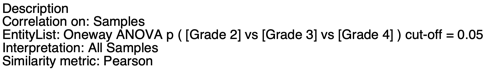

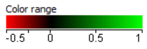

**Grade 2**

**Grade 3**

**Grade 4**

**Figure 3.** Transcriptional profiles of OA patients from different grades. (a) The hierarchal clustering and (b) correlation analysis shows variations in gene expression among osteoarthritis patients of different grades: Grade 2 (n=3), Grade 3 (n=3), and Grade 4 (n=4).

(a)Green denotes upregulated genes, red indicates downregulated genes, and black represents genes with stable expression. (b) Green shows positively correlated samples, red shows negatively correlated samples, and black shows samples not related.

**Table 3.** Upregulated and downregulated pathways and their associated genes from Grade 3 v/s Grade 2 comparison using GSEA database.

| **S.No.** | **Pathways**  **(Grade 3 vs Grade 2)** | **Category** | **Enrichment score** | **Count** | **FDR** | **Regulation** | **Genes** |
| --- | --- | --- | --- | --- | --- | --- | --- |
| 1 | Neutrophil degranulation | Reactome | 0.55 | 8 | 0.084 | Up | *ARPC5, SLC2A5, MME, PKP1, SERPINA3, ARG1, PTPRN2, CFD* |
| 2 | Developmental biology | Reactome | 0.24 | 20 | 0.095 | Up | *ARPC5, SEMA6D, MYF6, KALRN, RARB, HELZ2, KRTAP23-1, KRTAP4-3, LPL, NKX6-1, CACNA1C, LCE2D, PKP1, FOXF1, MYB, FOXA2, KRTAP10-4, SPRR2A, LCE1E, KCNQ3* |
| 3 | Process utilizing autophagic mechanism | BP | -0.38 | 11 | 0.069 | Down | *SVIP, DNM1L, FOXK1, CDK5, MAPK15, BAG3, RNF185, WDR45, CERS1, HSPA8, EEF1A2* |
| 4 | Positive regulation of programmed cell death | BP | -0.28 | 13 | 0.053 | Down | *PNMA1, GSN, DNM1L, STK4, OLFM1, CASP8, CDK5, STYXL1, IP6K2, NTSR1, EEF1A2, FAS, LTA* |
| 5 | Regulation of autophagy | BP | -0.41 | 9 | 0.071 | Down | *SVIP, DNM1L, FOXK1, CDK5, MAPK15, BAG3, WDR45, CERS1, EEF1A2* |
| 6 | Negative regulation of RNA biosynthetic process | BP | -0.17 | 27 | 0.087 | Down | *MAGEC2, FOXG1, ZC3H6, SRY, ZNF281, NRG1, FOXK1, ZNF343, TDG, CDK5, KDM5B, MAFK, TNFSF4, BAG3, MAGEF1, TCFL5, BTAF1, ZNF3, KAT6B, LRRFIP1, ZBTB26, ZNF589, HSPA8, SRF, SLA2, ZNF541, ZNF77* |
| 7 | RNA processing | BP | -0.18 | 18 | 0.083 | Down | *PRPF3, WDR33, SNORA8, SNORD22, PTCD2, SNORD36A, SNHG7, IWS1, SNORD86, SNORA66, CLK2, CCNL2, RSRP1, SNORD54, RRP12, HSPA8, SNORA3A, RPS21* |
| 8 | Negative regulation of nucleobase containing compound metabolic process | BP | -0.15 | 24 | 0.087 | Down | *ZNF281, NRG1, FOXK1, ZNF343, TDG, CDK5, KDM5B, MAFK, TNFSF4, BAG3, MAGEF1, TCFL5, BTAF1, ZNF3, KAT6B, LRRFIP1, ZBTB26, ZNF589, CERS1, HSPA8, SRF, SLA2, ZNF541, ZNF77* |
| 9 | Cell substrate adhesion | BP | -0.3 | 8 | 0.092 | Down | *MUC4, STK4, CDK5, EPHB3, LIMS2, SRF, FBLN2, MSLN* |
| 10 | Inflammatory response | BP | -0.21 | 12 | 0.088 | Down | *PNMA1, CNTF, RABGEF1, TNFSF4, PTGER3, LDLR, NFKBIB, HSPA8, CHST1, IL5RA, LTA, IL1R2* |
| 11 | Nucleolus | CC | -0.19 | 18 | 0.062 | Down | *THAP2, PNMA1, WDR33, SDHA, SNORA8, SNORD22, SRP19, SNORD36A, SNHG7, RABGEF1, SNORD86, SNORA66, KLF6, IP6K2, SNORD54, RRP12, HSPA8, SNORA3A* |
| 12 | Golgi apparatus | CC | -0.16 | 24 | 0.054 | Down | *FUT5, ICA1L, SUN5, TXNDC8, MUC4, KERA, MMP16, SVIP, ENTPD7, DNM1L, LPCAT1, MAPK15, YIF1B, PERP, FGF7, ST3GAL4, LDLR, NTSR1, PHETA1, CHST1, HS6ST2, BAIAP3, SLA2, MSLN* |
| 13 | Guanyl nucleotide binding | MF | -0.35 | 25 | 0.032 | Down | *RAB3A, PDE11A, NMUR2, RAB25, HHAT, EIF2S3B, GBP6, SEPTIN4, GFM1, MPPED2, RAB43, GIMD1, RAB41, RAB3C, SEPTIN14, RASEF, SRL, CNGA2, MMAA, RERGL, GBP7, DNM1L, RAB28, EEF1A2, RRAD* |
| 14 | Transcription regulator activity | MF | -0.14 | 30 | 0.019 | Down | *SRY, ZNF627, ZNF281, ZNF396, NRG1, ZBTB43, ZNF888, ZSCAN5C, FOXK1, ZNF343, TDG, KDM5B, ZNF319, MAFK, TCFL5, HLX, KLF6, BTAF1, ZNF833P, ZNF3, KAT6B, LRRFIP1, NFKBIB, ZBTB26, ZNF589, ZNF780A, SRF, ZNF222, ZNF541, ZNF77* |
| 15 | DNA binding transcription repressor activity | MF | -0.27 | 10 | 0.051 | Down | *ZNF281, FOXK1, ZNF343, MAFK, TCFL5, ZNF3, LRRFIP1, ZBTB26, ZNF589, ZNF77* |
| 16 | Cytokine signaling in immune system | Reactome | -0.27 | 14 | 0.004 | Down | *CNTF, CASP8, FAAP20, CLCF1, SOCS2, TNFSF4, IP6K2, NFKBIB, HSPA8, SLA2, IL5RA, LTA, IL1R2* |
| 17 | Transcriptional regulation by TP53 | Reactome | -0.5 | 14 | 0.003 | Down | *TAF1, ELOA2, RAD9B, TAF7L, MSH2, PRKAA2, FANCC, TP53AIP1, NUAK1, CASP10, CDK5, PERP, FAS* |
| 18 | RNA polymerase II transcription | Reactome | -0.21 | 19 | 0.007 | Down | *ZNF705G, NR4A1, PSMB7, FOXG1, ZNF627, WDR33, CASP10, ZNF343, CDK5, KDM5B, IWS1, PERP, ZNF3, ZNF589, SRF, ZNF222, FAS, ZNF77* |

*False Discovery Rate (FDR)≤0.1, BP-Biological pathway, CC-Cellular component, MF-Molecular function

**Table 4.** Upregulated and downregulated pathways and their associated genes from Grade 4 v/s Grade 2 comparison using GSEA database.

| **S.No.** | **Pathways (Grade 4 vs Grade 2)** | **Enrichment Score** | **Category** | **Count** | **FDR** | **Regulation** | **Genes** |
| --- | --- | --- | --- | --- | --- | --- | --- |
| 1 | Defense response to symbiont | 0.61 | BP | 17 | 0.051 | Up | *STAT1, RSAD2, IFIT3, GBP1, USP18, IFIT1, HERC5, DDX60, IFIT2, IL21, RTP4, PARP9, IFIH1, DDX60L, OAS2, DHX58, USP44* |
| 2 | Response to virus | 0.56 | BP | 18 | 0.027 | Up | *STAT1, RSAD2, IFIT3, GBP1, USP18, IFIT1, IFI44, HERC5, DDX60, IFIT2, IL21, RTP4, PARP9, IFIH1, DDX60L, OAS2, DHX58, USP44* |
| 3 | Response to organic cyclic compound | 0.61 | BP | 7 | 0.018 | Up | *CSN1S1, STAT1, USP18, SLC1A3, NR4A3, IFIT1, CYP27B1* |
| 4 | Biological process involved in interspecies interaction between organisms | 0.46 | BP | 29 | 0.02 | Up | *NEURL3, STAT1, RSAD2, IFIT3, GBP1, USP18, IL31RA, IFIT1, IFI44, CMPK2, HERC5, CYP27B1, DDX60, IFIT2, PDCD1LG2, IL21, RTP4, RNF213, LEAP2, PARP9, BATF2, IFIH1, DDX60L, TAP2, OAS2, CX3CL1, TFRC, DHX58, USP44* |
| 5 | Defense response to other organism | 0.49 | BP | 22 | 0.023 | Up | *NEURL3, STAT1, RSAD2, IFIT3, GBP1, USP18, IL31RA, IFIT1, HERC5, CYP27B1, DDX60, IFIT2, IL21, RTP4, RNF213, LEAP2, PARP9, BATF2, IFIH1, DDX60L, OAS2, CX3CL1* |
| 6 | Innate immune response | 0.53 | BP | 12 | 0.026 | Up | *NEURL3, STAT1, RSAD2, IFIT3, GBP1, USP18, IFIT1, HERC5, CYP27B1, DDX60, IFIT2, IL21* |
| 7 | Response to hormone | 0.59 | BP | 5 | 0.064 | Up | *CSN1S1, STAT1, RARB, NR4A3, CYP27B1* |
| 8 | External encapsulating structure | 0.53 | CC | 4 | 0.092 | Up | *AMTN, ADAMDEC1, LAMC2, CASK* |
| 9 | Interferon signaling | 0.71 | Reactome | 10 | 0.00001 | Up | *USP41, STAT1, RSAD2, IFIT3, GBP1, USP18, IFIT1, HERC5, XAF1, IFIT2* |
| 10 | Cytokine signaling in immune system | 0.54 | Reactome | 14 | 0.004 | Up | *IL20, USP41, STAT1, RSAD2, IFIT3, GBP1, USP18, IL31RA, IFIT1, HERC5, XAF1, IFIT2, IL21, DUSP4* |
| 11 | Transport of small molecules | 0.54 | Reactome | 4 | 0.037 | Up | *CSN1S1, SLC1A3, CLCNKA, SLC30A2* |
| 12 | Supramolecular polymer | -0.57 | CC | 13 | 0.002 | Down | *DES, ARL3, ACTA2, TMOD3, MAP4, BAG3, MAP1LC3C, PKP1, DST, KRTAP10-4, SVIL, NES, TPPP3* |
| 13 | Supramolecular complex | -0.45 | CC | 21 | 0.007 | Down | *CSRP1, COL7A1, MOV10, CNOT2, GIGYF2, ARHGAP6, DES, ARL3, ACTA2, TMOD3, MAP4, BAG3, MAP1LC3C, PKP1, DST, KRTAP10-4, SVIL, CENPM, NES, TPPP3, TDRD1* |
| 14 | Microtubule cytoskeleton | -0.38 | CC | 16 | 0.054 | Down | *NGRN, MAPK15, AHI1, ARL3, HECW2, MAP4, MAP1LC3C, DST, OLA1, SVIL, CDH23, CEP126, DTX4, SPEF2, TPPP3, BRSK1* |
| 15 | Golgi apparatus | -0.34 | CC | 7 | 0.055 | Down | *FGF7, HS6ST2, BAIAP3, SPEF2, YIF1B, CHST1, MSLN* |

*False Discovery Rate (FDR)≤0.1, BP-Biological pathway, CC-Cellular component, MF-Molecular function

**Table 5.** Upregulated and downregulated pathways and their associated genes from Grade 4 v/s Grade 3 comparison using GSEA database.

| **S.No.** | **Pathways**  **(Grade 4 vs Grade 3)** | **Enrichment score** | **Category** | **Count** | **FDR** | **Regulation** | **Genes** |
| --- | --- | --- | --- | --- | --- | --- | --- |
| 1 | Response to virus | 0.68 | BP | 23 | 0.0001 | Up | *GBP1, STAT1, RSAD2, DDX60, USP18, IFIT3, HSPA8, PARP9,RTP4, MOV10, IFIT2, RNF185, HERC5, ACTA2, USP44, IFIT1, TNFSF4, IFI44, NT5C3A, DHX58, IFIH1, OAS2, DDX60L* |
| 2 | Biological process involved in interspecies interaction between organisms | 0.42 | BP | 45 | 0.0001 | Up | *GBP1, LTA, NEURL3, STAT1, RSAD2, CMPK2, DDX60, USP18, IFIT3, HSPA8, IL31RA, PARP9, RTP4, MMRN2, RNF213, LDLR, TFRC, MOV10, CYP27B1, IFIT2, RNF185, HERC5, ACTA2, USP44, IFIT1, TNFSF4, OTUD4, WDFY1, IFI44, TAP2, CASP8, NT5C3A, OCIAD2, NFKBIB, DHX58, SMC5, IFIH1, OAS2, DDX60L, WRNIP1, CLEC4A, LEAP2, PDCD1LG2, ZC3HAV1, F2* |
| 3 | Defense response to symbiont | 0.72 | BP | 20 | 0.0001 | Up | *GBP1, STAT1, RSAD2, DDX60, USP18, IFIT3, HSPA8, PARP9, RTP4, MOV10, IFIT2, RNF185, HERC5, USP44, IFIT1, NT5C3A, DHX58, IFIH1, OAS2, DDX60L* |
| 4 | Regulation of immune system process | 0.49 | BP | 41 | 0.0001 | Up | *GBP1, LTA, SLA2, STAT1, RSAD2, ZBTB26, DDX60, USP18, IL20, HSPA8, IL31RA, PARP9, LDLR, NR4A3, HLX, TFRC, IL15RA, GPR68, RNF185, TNFSF4, OTUD4, WDFY1, TAP2, CASP8, RABGEF1, ST3GAL4, DNM1L, DHX58, IFIH1, RC3H2, ZBTB25* |
| 5 | Positive regulation of immune response | 0.6 | BP | 19 | 0.0001 | Up | *GBP1, LTA, SLA2, RSAD2, DDX60, HSPA8, PARP9, NR4A3, HLX, TFRC, RNF185, TNFSF4, OTUD4, WDFY1, TAP2, RABGEF1, DHX58, IFIH1, RC3H2* |
| 6 | Regulation of immune response | 0.54 | BP | 21 | 0.0001 | Up | *GBP1, LTA, SLA2, RSAD2, DDX60, USP18, HSPA8, PARP9, NR4A3, HLX, TFRC, RNF185, TNFSF4, OTUD4, WDFY1, TAP2, CASP8, RABGEF1, DHX58, IFIH1, RC3H2* |
| 7 | Defense response to other organism | 0.43 | BP | 35 | 0.0001 | Up | *GBP1, LTA, NEURL3, STAT1, RSAD2, DDX60, USP18, IFIT3, HSPA8, IL31RA, PARP9, RTP4, MMRN2, RNF213, MOV10, CYP27B1, IFIT2, RNF185, HERC5, USP44, IFIT1, TNFSF4, OTUD4, WDFY1, CASP8, NT5C3A, DHX58, IFIH1, OAS2, DDX60L, WRNIP1, CLEC4A, LEAP2, ZC3HAV1, F2* |
| 8 | Positive regulation of immune system process | 0.53 | BP | 24 | 0.0001 | Up | *GBP1, LTA, SLA2, RSAD2, DDX60, IL20, HSPA8, PARP9, NR4A3, HLX, TFRC, IL15RA, GPR68, RNF185, TNFSF4, OTUD4, WDFY1, TAP2, CASP8, RABGEF1, DNM1L, DHX58, IFIH1, RC3H2* |
| 9 | Defense response | 0.37 | BP | 41 | 0.0001 | Up | *GBP1, LTA, NEURL3, STAT1, RSAD2, DDX60, USP18, IFIT3, IL20, HSPA8, IL31RA, PARP9, RTP4, MMRN2, RNF213, LDLR, TFRC, MOV10, GPR68, CYP27B1, IFIT2, RNF185, HERC5, USP44, IFIT1, TNFSF4, OTUD4, WDFY1, CASP8, RABGEF1, NT5C3A, NFKBIB, DHX58, PNMA1, IFIH1, OAS2, DDX60L, DLEC1, FANCA, WRNIP1, CLEC4A* |
| 10 | Viral process | 0.73 | BP | 14 | 0.0001 | Up | *STAT1, RSAD2, HSPA8, PARP9, LDLR, TFRC, IFIT1, ST3GAL4, SMC5, IFIH1, OAS2, MVB12A, FAM111A, ZC3HAV1* |
| 11 | Regulation of defense response | 0.47 | BP | 20 | 0.0001 | Up | *LTA, STAT1, RSAD2, DDX60, USP18, IL20, HSPA8, PARP9, MMRN2, LDLR, RNF185, HERC5, TNFSF4, OTUD4, WDFY1, CASP8, RABGEF1, DHX58, IFIH1, FANCA* |
| 12 | Hemopoiesis | 0.51 | BP | 20 | 0.0001 | Up | *STAT1, RSAD2, IL20, KLF6, SNRK, IL31RA, SRF, HLX, TFRC, IL15RA, GPR68, MMP21, USP44, TNFSF4, CASP8, STK4, RC3H2, FANCA, CARTPT, DACT2* |
| 13 | Immune response regulating signaling pathway | 0.66 | BP | 12 | 0.0001 | Up | *GBP1, SLA2, RSAD2, DDX60, HSPA8, NR4A3, OTUD4, WDFY1, RABGEF1, DHX58, IFIH1, RC3H2* |
| 14 | Innate immune response | 0.4 | BP | 23 | 0.0001 | Up | *GBP1, NEURL3, STAT1, RSAD2, DDX60, USP18, IFIT3, HSPA8, PARP9, CYP27B1, IFIT2, RNF185, HERC5, USP44, IFIT1, OTUD4, WDFY1, CASP8, DHX58, IFIH1, OAS2, WRNIP1, CLEC4A* |
| 15 | Regulation of response to biotic stimulus | 0.52 | BP | 14 | 0.0001 | Up | *STAT1, RSAD2, DDX60, USP18, HSPA8, PARP9, MMRN2, RNF185, HERC5, OTUD4, WDFY1, CASP8, DHX58, IFIH1* |
| 16 | DNA metabolic process | 0.44 | BP | 23 | 0.0001 | Up | *DNASE2B, PARP9, TFRC, ASF1A, TDG, WDR33, USP44, TNFSF4, OTUD4, DFFB, ESCO1, MAGEF1, SMC5, RAD17, RRM2, FANCA, WRNIP1, SMARCAD1, DPF1, TAF5, PMS1, NSD2, FAM111A* |
| 17 | Cytokine production | 0.46 | BP | 18 | 0.0001 | Up | *GBP1, LTA, STAT1, RSAD2, ZBTB26, XAF1, NR4A3, TNFSF4, CASP8, RABGEF1, DHX58, IFIH1, ZBTB25, OAS2, CLEC4A, PDCD1LG2, ZC3HAV1, F2* |
| 18 | Positive regulation of programmed cell death | 0.51 | BP | 12 | 0.0001 | Up | *GBP1, LTA, NTSR1, FAS, IP6K2, CDK5, IFIT2, CASP8, STYXL1, STK4, DNM1L, PNMA1* |
| 19 | Positive regulation of defense response | 0.54 | BP | 13 | 0.0001 | Up | *LTA, RSAD2, DDX60, HSPA8, PARP9, MMRN2, LDLR, RNF185, TNFSF4, OTUD4, WDFY1, DHX58, IFIH1* |
| 20 | Activation of immune response | 0.54 | BP | 12 | 0.0001 | Up | *GBP1, SLA2, RSAD2, DDX60, HSPA8, NR4A3, OTUD4, WDFY1, RABGEF1, DHX58, IFIH1, RC3H2* |
| 21 | Nucleolus | 0.46 | CC | 34 | 0.0001 | Up | *STAT1, RSAD2, SNORA3A, HSPA8, KLF6, SNORA66, RNF213, SNORD54, CASK, IP6K2, WDR33, SNORD22, DFFB, WDFY1, FAM9A, PDK3, RABGEF1, SNORA8, SDHA, SNORD36A, SNORD36B, PNMA1, SRP19, RAD17, C1orf131, POLR1D, COX7A2L, SNORD86, RRP12, TAF5, NSD2, SNHG7, RRN3, FAM111A* |
| 22 | Coated vesicle | 0.6 | CC | 11 | 0.0001 | Up | *HSPA8, LDLR, TFRC, ENTHD1, STX17, EPN1, DDHD2, COL7A1, MCFD2, ATP6V1B2, SCYL2* |
| 23 | Mitochondrion | 0.33 | CC | 31 | 0.0001 | Up | *RSAD2, CMPK2, IFIT3, XAF1, PARP9, PTCD2, SLC25A45, CYP27B1, SAMD9L, RNF185, GCDH, PCCB, AMT, LDHB, PDK3, CASP8, STYXL1, SIRT5, ISCA1, MRPS5, STX17, SDHA, DNM1L, PERP, ACSS1, IFIH1, COX7A2L, ACACB, DARS2, CAPRIN2, SFXN4* |
| 24 | Nuclear body | 0.37 | CC | 17 | 0.004 | Up | *FAS, TDG, CCNL2, PARP11, CLK2, TAP2, NT5C3A, STK4, SRP19, SMC5, FAM76B, PRPF3, USPL1, ZNF451, SIMC1, SUGP2, RNF6* |
| 25 | Catalytic complex | 0.27 | CC | 20 | 0.029 | Up | *SLA2, ZNF541, DNHD1, KDM3A, CDK5, CUL1, CCNL2, PCCB, LDHB, CASP8, SDHA, MAT2B, SMC5, RRM2, ATP6V1B2, POLR1D, COX7A2L, PCGF3, DPF1, TAF5* |
| 26 | Endosome | 0.3 | CC | 13 | 0.026 | Up | *SLA2, NEURL3, HSPA8, STAMBPL1, LDLR, TFRC, IL15RA, SAMD9L, ENTHD1, WDFY1, RABGEF1, OCIAD2, EPN1* |
| 27 | Mitochondrial matrix | 0.42 | CC | 10 | 0.03 | Up | *GCDH, PCCB, AMT, PDK3, STYXL1, SIRT5, ISCA1, MRPS5, ACSS1, DARS2* |
| 28 | Envelope | 0.27 | CC | 21 | 0.029 | Up | *RSAD2, SLC25A45, IL15RA, CYP27B1, RNF185, PARP11, LDHB, CASP8, SIRT5, MRPS5, FAM156B, SDHA, DNM1L, COX7A2L, QSOX2, RRP12, ACACB, NUP58, PARP8, SFXN4, RNF6* |
| 29 | Golgi apparatus | 0.24 | CC | 25 | 0.031 | Up | *GBP1, SLA2, NTSR1, RSAD2, LDLR, IL15RA, PLEKHA8, WDFY1, LPCAT1, AGRN, OCIAD2, ST3GAL4, DNM1L, PERP, DDHD2, MCFD2, QSOX2, SCYL2, F2, SGSM1, MYMK, PICK1, ARL3, LRP1, RNF183* |
| 30 | Side of membrane | 0.28 | CC | 8 | 0.041 | Up | *STAC, NTSR1, FAS, HSPA8, IL31RA, PKP4, LDLR, TFRC* |
| 31 | Vesicle membrane | 0.23 | CC | 25 | 0.067 | Up | *GBP1, SLA2, HSPA8, LDLR, TFRC, IL15RA, ENTHD1, TAP2, LPCAT1, RABGEF1, STX17, DNM1L, EPN1, MVB12A, MCFD2, ATP6V1B2, SCYL2, SGSM1, SLC5A7, PICK1, DMBT1, LRP1, ATP13A4, WNT4, AQP7* |
| 32 | Mitochondrial envelope | 0.29 | CC | 15 | 0.088 | Up | *RSAD2, SLC25A45, CYP27B1, RNF185, LDHB, CASP8, SIRT5, MRPS5, SDHA, DNM1L, COX7A2L, ACACB, SFXN4, STARD13, PRODH2* |
| 33 | Catalytic activity acting on a nucleic acid | 0.62 | MF | 17 | 0.0001 | Up | *DDX60, DNASE2B, MOV10, TDG, DQX1, BTAF1, DFFB, NT5C3A, DHX58, RAD17, IFIH1, DDX60L, POLR1D, WRNIP1, SMARCAD1, PMS1, DARS2* |
| 34 | Purine nucleotide binding | 0.35 | MF | 48 | 0.0001 | Up | *GBP1, CMPK2, DDX60, HSPA8, DNHD1, SNRK, PARP9, RNF213, MOV10, CASK, IP6K2, CDK5, RAB28, TDG, GIMAP4, PASK, GCDH, FMO5, ACTA2, DQX1, PCCB, BTAF1, CLK2, TAP2, LDHB, PDK3, STK31, SIRT5, STK4, DNM1L, DHX58, SMC5, ACSS1, RAD17, IFIH1, OAS2, DDX60L, MAST4, ATP6V1B2, WRNIP1, RRAD, ACACB, SCYL2, SMARCAD1, PMS1, DARS2, UBA3, NEK3* |
| 35 | Adenyl nucleotide binding | 0.34 | MF | 44 | 0.0001 | Up | *CMPK2, DDX60, HSPA8, DNHD1, SNRK, PARP9, RNF213, MOV10, CASK, IP6K2, CDK5, TDG, PASK, GCDH, FMO5, ACTA2, DQX1, PCCB, BTAF1, CLK2, TAP2, LDHB, PDK3, STK31, SIRT5, STK4, DHX58, SMC5, ACSS1, RAD17, IFIH1, OAS2, DDX60L, MAST4, ATP6V1B2, WRNIP1, ACACB, SCYL2, SMARCAD1, PMS1, DARS2, UBA3, NEK3, ZC3HAV1* |
| 36 | Hydrolase activity acting on acid anhydrides | 0.41 | MF | 18 | 0.0001 | Up | *GBP1, DDX60, HSPA8, RNF213, MOV10, RAB28, DQX1, BTAF1, TAP2, DNM1L, DHX58, SMC5, IFIH1, DDX60L, WRNIP1, RRAD, SMARCAD1, PMS1* |
| 37 | Transcription regulator activity | 0.28 | MF | 32 | 0.0001 | Up | *ZNF77, ZNF541, STAT1, ZBTB26, ZNF780A, HESX1, KLF6, ZNF69, IL31RA, KDM3A, PARP9, NR4A3, SRF, HLX, ZNF589, TDG, FOXK1, ZSCAN5C, BTAF1, ZNF222, ZNF519, ZNF281, ZNF3, NFKBIB, TSTD1, ZNF496, ZBTB43, NRF1, ZNF319, ZBTB25, LRRFIP1, PTF1A* |
| 38 | ATP dependent activity | 0.4 | MF | 18 | 0.001 | Up | *DDX60, HSPA8, DNHD1, RNF213, MOV10, DQX1, BTAF1, TAP2, DHX58, SMC5, RAD17, IFIH1, DDX60L, ATP6V1B2, WRNIP1, SMARCAD1, PMS1* |
| 39 | ATP hydrolysis activity | 0.46 | MF | 14 | 0.001 | Up | *DDX60, HSPA8, RNF213, MOV10, DQX1, BTAF1, TAP2, DHX58, SMC5, IFIH1, DDX60L, WRNIP1, SMARCAD1, PMS1* |
| 40 | DNA binding transcription factor activity | 0.26 | MF | 28 | 0.007 | Up | *ZNF77, STAT1, ZBTB26, ZNF780A, HESX1, KLF6, ZNF69, NR4A3, SRF, HLX, ZNF589, FOXK1, ZSCAN5C, ZNF222, ZNF519, ZNF281, ZNF3, TSTD1, ZNF496, ZBTB43, NRF1, ZNF319, ZBTB25, LRRFIP1, PTF1A, ZNF684, ZNF451, ZNF24* |
| 41 | DNA binding transcription repressor activity | 0.43 | MF | 11 | 0.008 | Up | *ZNF77, ZBTB26, HESX1, ZNF69, ZNF589, FOXK1, ZNF281, ZNF3, ZBTB25, LRRFIP1, ZNF684* |
| 42 | Molecular adaptor activity | 0.46 | MF | 8 | 0.009 | Up | *SLA2, USP18, HSPA8, LDLR, CUL1, OTUD4, STX17, KLHL17* |
| 43 | Zinc ion binding | 0.3 | MF | 17 | 0.01 | Up | *ADAMDEC1, XAF1, CA2, RABIF, NR4A3, CPXM1, CPA6, MMP21, USP44, WDFY1, ESCO1, CZIB, RABGEF1, SIRT5, ZNF3, DHX58, IFIH1* |
| 44 | Sequence specific DNA binding | 0.24 | MF | 30 | 0.016 | Up | *ZNF77, STAT1, ZBTB26, ZNF780A, HESX1, KLF6, ZNF69, NR4A3, SRF, HLX, ZNF589, FOXK1, ZSCAN5C, ZNF519, ZNF281, ZNF3, TSTD1, ZNF496, ZBTB43, NRF1, ZNF319, ZBTB25, LRRFIP1, PTF1A, ZNF684, ZNF451, DPF1, ZNF24, NSD2, RRN3* |
| 45 | Transition metal ion binding | 0.25 | MF | 20 | 0.02 | Up | *ADAMDEC1, XAF1, CA2, RABIF, KDM3A, NR4A3, CPXM1, CPA6, CYP27B1, MMP21, USP44, WDFY1, ESCO1, CZIB, RABGEF1, SIRT5, ZNF3, DHX58, RRM2, IFIH1* |
| 46 | Transcription factor binding | 0.38 | MF | 12 | 0.023 | Up | *STAT1, KDM3A, PARP9, NR4A3, SRF, TDG, BTAF1, STK4, ZBTB43, BDP1, RNF6, DACT2* |
| 47 | Transferase activity transferring phosphorus containing groups | 0.28 | MF | 19 | 0.029 | Up | *CMPK2, SNRK, PARP9, CASK, IP6K2, CDK5, PARP11, PASK, CLK2, PDK3, STK31, NT5C3A, STK4, OAS2, MAST4, POLR1D, SCYL2, PARP8, NEK3* |
| 48 | Protein homodimerization activity | 0.31 | MF | 12 | 0.033 | Up | *GBP1, STAT1, HESX1, NR4A3, SRF, TFRC, ADRB3, STK4, DNM1L, NRF1, RRM2, LRRFIP1* |
| 49 | DNA binding transcription factor binding | 0.4 | MF | 7 | 0.037 | Up | *STAT1, KDM3A, PARP9, NR4A3, SRF, TDG, STK4* |
| 50 | Enzyme regulator activity | 0.25 | MF | 25 | 0.042 | Up | *SLA2, PPME1, RABIF, PARP9, ARHGAP19, CCNL2, RASA4, STYXL1, RABGEF1, STK4, DNM1L, MAT2B, COL7A1, CALML4, WRNIP1, SIMC1, CPEB2, EPS8L1, PARP8, SGSM1, STARD13, PAPLN, ARHGEF25, ARHGAP29, ADAP2* |
| 51 | Cis regulatory region sequence specific dna binding | 0.22 | MF | 18 | 0.053 | Up | *STAT1, ZBTB26, ZNF780A, HESX1, KLF6, NR4A3, SRF, FOXK1, ZSCAN5C, ZNF519, ZNF281, ZNF496, ZBTB43, NRF1, ZNF319, ZBTB25, LRRFIP1, PTF1A* |
| 52 | Peptidase activity | 0.28 | MF | 10 | 0.069 | Up | *ADAMDEC1, CAPN12, USP18, STAMBPL1, CPXM1, CPA6, MMP21, USP44, OTUD4, CASP8* |
| 53 | Cytokine signaling in immune system | 0.6 | Reactome | 26 | 0.0001 | Up | *USP41, GBP1, LTA, SLA2, STAT1, RSAD2, USP18, IFIT3, IL20, XAF1, HSPA8, IL31RA, IL15RA, IP6K2, CUL1, IFIT2, HERC5, IFIT1, TNFSF4, CASP8, NFKBIB, OAS2, FANCA, UBA3, NUP58, DUSP4* |
| 54 | Interferon signaling | 0.79 | Reactome | 15 | 0.0001 | Up | *USP41, GBP1, STAT1, RSAD2, USP18, IFIT3, XAF1, HSPA8, IP6K2, IFIT2, HERC5, IFIT1, OAS2, FANCA, NUP58* |
| 55 | RNA polymerase II transcription | 0.44 | Reactome | 21 | 0.0001 | Up | *ZNF77, STAT1, FAS, NR4A3, SRF, MOV10, CDK5, ZNF589, CUL1, WDR33, ZNF222, ZNF519, PERP, ZNF3, ZNF496, RAD17, RRM2, IWS1, ZNF684, COX7A2L, TAF5* |
| 56 | Signaling by interleukins | 0.47 | Reactome | 9 | 0.001 | Up | *STAT1, USP18, IL20, HSPA8, IL31RA, IL15RA, CUL1, CASP8, NFKBIB* |
| 57 | Adaptive immune system | 0.4 | Reactome | 12 | 0.008 | Up | *SIGLEC12, RNF213, CUL1, HERC5, TAP2, NFKBIB, CLEC2D, UBA3, UBE3B, PDCD1LG2, RNF6, TRIM71* |
| 58 | Post translational protein modification | 0.23 | Reactome | 20 | 0.043 | Up | *USP18, HSPA8, STAMBPL1, TDG, CUL1, RNF185, FOXK1, USP44, PIGB, ST3GAL4, STX17, MAT2B, SMC5, IFIH1, COL7A1, MCFD2, AMTN, UBA3, NUP58, F2* |
| 59 | Actin filament organization | -0.59 | BP | 14 | 0.009 | Down | *FMN1, PFN3, SVIL, POF1B, STMN1, TLE6, ARHGAP6, WASHC1, RHOJ, WASF3, CCL21, PPM1E, MAGEL2, ARPC5* |
| 60 | Regulation of actin filament based process | -0.59 | BP | 13 | 0.012 | Down | *FMN1, PFN3, SVIL, STMN1, ARHGAP6, STC1, WASHC1, CACNA1C, WASF3, CCL21, PPM1E, MAGEL2, ARPC5* |
| 61 | Supramolecular fiber organization | -0.47 | BP | 19 | 0.011 | Down | *FMN1, KRT20, PFN3, SVIL, POF1B, DES, STMN1, TLE6, ARHGAP6, IAPP, MAP4, WASHC1, RHOJ, WASF3, CCL21, PPM1E, PKP1, MAGEL2, ARPC5* |
| 62 | Regulation of organelle organization | -0.43 | BP | 21 | 0.017 | Down | *CEP295NL, FMN1, C10orf90, PFN3, SVIL, FZD9, HECW2, STMN1, ARHGAP6, IAPP, MAP4, NAF1, CPLX2, WASHC1, NES, WASF3, PDE2A, CCL21, PPM1E, MAGEL2, ARPC5* |
| 63 | Regulation of cytoskeleton organization | -0.54 | BP | 12 | 0.014 | Down | *FMN1, PFN3, SVIL, STMN1, ARHGAP6, WASHC1, NES, WASF3, CCL21, PPM1E, MAGEL2, ARPC5* |
| 64 | Regulation of supramolecular fiber organization | -0.53 | BP | 12 | 0.025 | Down | *FMN1, PFN3, SVIL, STMN1, ARHGAP6, IAPP, WASHC1, WASF3, CCL21, PPM1E, MAGEL2, ARPC5* |
| 65 | Actin filament based process | -0.42 | BP | 18 | 0.031 | Down | *FMN1, KCNN2, PFN3, SVIL, POF1B, STMN1, TLE6, ARHGAP6, STC1, KCNH2, WASHC1, RHOJ, CACNA1C, WASF3, CCL21, PPM1E, MAGEL2, ARPC5* |
| 66 | Supramolecular polymer | -0.41 | CC | 21 | 0.035 | Down | *FMN1, KRT20, KCNN2, LRRC27, SVIL, POF1B, DES, STMN1, ARHGAP6, MAP4, NES, KRTAP23-1, CACNA1C, DST, MYZAP, KRTAP4-3, LMNTD1, C10orf71, MAP1LC3C, KRTAP10-4, PKP1* |
| 67 | Polymeric cytoskeletal fiber | -0.45 | CC | 16 | 0.024 | Down | *FMN1, KRT20, SVIL, POF1B, DES, STMN1, ARHGAP6, MAP4, NES, KRTAP23-1, DST, KRTAP4-3, LMNTD1, MAP1LC3C, KRTAP10-4, PKP1* |
| 68 | Supramolecular complex | -0.36 | CC | 26 | 0.02 | Down | *FMN1, KRT20, KCNN2, HIPK2, LRRC27, SVIL, POF1B, DES, HELZ2, STMN1, RBPMS, ARHGAP6, GIGYF2, MAP4, NES, KRTAP23-1, CACNA1C, DST, MYZAP, KRTAP4-3, LMNTD1, C10orf71, MAP1LC3C, TDRD1, KRTAP10-4, PKP1* |
| 69 | Anchoring junction | -0.37 | CC | 18 | 0.021 | Down | *SVIL, POF1B, PANX3, DES, ILDR2, JAK1, AHI1, CDH3, GRHL2, CLDN4, VSIG10L2, DST, MYZAP, CCDC85A, GJA3, MME, PKP1, ARPC5* |
| 70 | Secretory granule | -0.38 | CC | 5 | 0.077 | Down | *RAB44, CADPS, MME, PKP1, ARPC5* |
| 71 | Secretory vesicle | -0.35 | CC | 10 | 0.082 | Down | *TMPRSS12, CFD, ARG1, UNC13C, RAB44, CADPS, BRSK1, MME, PKP1, ARPC5* |
| 72 | Developmental biology | -0.34 | Reactome | 20 | 0.093 | Down | *SPRR2A, FOXF1, HELZ2, FOXA2, KCNQ3, FOXH1, IAPP, NKX6-1, KALRN, LCE2D, KRTAP23-1, CACNA1C, KRTAP4-3, FOXA3, RPS20, MYF6, KRTAP10-4, PKP1, SEMA6D, ARPC5* |

*False Discovery Rate (FDR)≤0.1, BP-Biological pathway, CC-Cellular component, MF-Molecular function

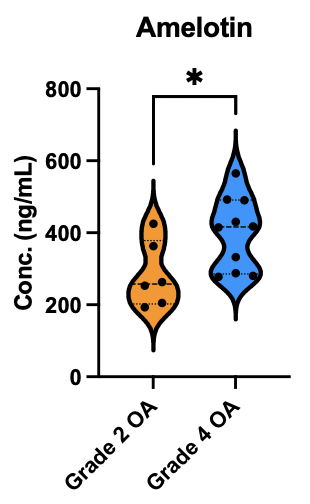

**Figure 4**. Amelotin protein quantification using ELISA in synovial fluid samples of late (grade 4) vs early (grade 2) osteoarthritis patients. *p value≤0.05
